# CRISPR/Cas9 screenings unearth protein arginine methyltransferase 7 as a novel driver of metastasis in prostate cancer

**DOI:** 10.1101/2023.07.20.549704

**Authors:** Maria Rodrigo-Faus, Africa Vincelle-Nieto, Natalia Vidal, Javier Puente, Melchor Saiz-Pardo, Alejandra Lopez-Garcia, Marina Mendiburu-Eliçabe, Nerea Palao, Cristina Baquero, Angel M Cuesta, Hui-Qi Qu, Hakon Hakonarson, Monica Musteanu, Armando Reyes-Palomares, Almudena Porras, Paloma Bragado, Alvaro Gutierrez-Uzquiza

## Abstract

Owing to the inefficacy of available treatments, the survival rate of patients with metastatic prostate cancer (mPCa) is severely decreased. Therefore, it is crucial to identify new therapeutic targets to increase their survival. This study aim was to identify the most relevant regulators of mPCa onset by performing two high-throughput CRISPR/Cas9 screenings. Furthermore, some of the top hits were validated using small interfering RNA (siRNA) technology, with protein arginine methyltransferase 7 *(PRMT7)* being the best candidate. Its inhibition or depletion via CRISPR significantly reduced mPCa cell capacities *in vitro*. Moreover, *PRMT7* ablation reduced mPCa appearance in chicken chorioallantoic membrane and mouse xenograft assays. Molecularly, PRMT7 reprograms the expression of several adhesion molecules through methylation of several transcription factors, such as FoxK1 or NR1H2, which results in primary tumor PCa cell adhesion loss and motility gain. Importantly, *PRMT7* is upregulated in advanced stages of Spanish PCa tumor samples and PRMT7 pharmacological inhibition reduces the dissemination of mPCa cells. Thus, here is shown that *PRMT7* is a potential therapeutic target and biomarker of mPCa.

## INTRODUCTION

Metastasis is becoming a major challenge in cancer treatment. It is the main cause of death in cancer patients, especially prostate cancer (PCa) patients(1). PCa remains the second most frequent cancer and the fifth leading cause of cancer-related death among men(2). Owing to on- going improvements in tumor diagnostic technologies and treatments, patients who remain in a non-disseminated PCa stage of the disease have a good outcome. In contrast, the mean survival of patients with PCa who develop distant metastases is less than 3 years(3). These patients are commonly administered androgen-castration hormonal therapy along with conventional chemotherapy(James *et al*, 2016). Additional treatments are also available; however, these approaches are not sufficiently effective to manage metastatic PCa (mPCa) patients. Hence, there is an urgent need to find new targets and biomarkers to establish effective therapies or to better adjust them to improve mPCa patient survival.

Despite the alarming number of deaths from metastatic cancer each year, the occurrence of metastasis remains poorly understood. Some biological changes have been described as crucial in promoting localized primary tumor progression to a secondary foci dissemination stage. They involve primary tumor cell adhesion loss, degradation of the basement membrane, intravasation to the circulatory system, survival in the bloodstream, extravasation into distant organs, and generation of a new tumor bulk(6). For some types of cancer, primary tumor cells have a preferred niche for colonization, as highlighted by the Stephan Paget “seed-and-soil” hypothesis(7). This is the case for PCa tumor cells, in which approximately 70% of metastases are found in the bone marrow(8).

Metastatic tumor cells display high levels of genomic instability and harbor several epigenetic alterations that may empower them with the ability to disseminate and generate a new tumor burden. While the exact mPCa gene signature remains unknown, mPCa cancer cells frequently bear alterations in androgen receptor signaling(9), mutations in *TP53* and RB1 loss(10), PTEN loss, overactivated Akt signaling, *ETS* gene rearrangements, and deleterious mutations in some DNA-repair genes such as *BRCA2*(11). Moreover, some studies have also highlighted the importance of epigenetic factors such as *EZH2*, in which ectopic overexpression correlates with metastasis and poor prognosis in PCa patients(12). Nonetheless, much remains to be investigated to fully understand the mechanisms that enhance the migratory and invasive abilities of PCa tumor cells.

Some single-gene studies have unearthed mPCa regulators such as *WNT5A*(13), *MAP4K4*(14) and *PPP1CA*(15). However, given the complexity of the biological mechanisms implicated in metastasis onset, high-throughput screening assays seem more optimal for uncovering the most relevant regulators of this process. In this context, CRISPR/Cas9 screening methods are becoming increasingly powerful. This approach is based on CRISPR/Cas9 genome editing that makes it possible to selectively modify the genome(16). Shalem et al.(17) developed Cas9/sgRNAs libraries using lentiviruses that can facilitate both positive and negative loss-of-function (LOF) screenings in mammalian cells. *In vitro* and *in vivo* CRISPR screenings have been successfully performed during the last years in different cancer models, including leukemia and lung cancerCheng *et al*, 2021). Notwithstanding its usefulness, to the best of our knowledge, only two CRISPR/Cas9 library studies have been reported for PCa. One of study evaluated genes related to arginine deprivation therapy resistance(20) and another study examined metformin treatment resistance(21). Therefore, the aim of this study was to conduct high-throughput *in vitro* screening assays using the human GeCKO CRISPR/Cas9 library(17), to identify the most important genes and biological processes involved in metastasis onset in PCa patients with the purpose of establishing new actionable targets and/or biomarkers of mPCa.

Briefly, we identified the protein arginine methyltransferase 7 (*PRMT7*) as an essential gene in mPCa. Further investigation demonstrated that PRMT7 induces a switch in the expression of cellular adhesion molecules, at least in part through FoxK1, NR1H2 and/or NCOA2/3 transcription factor methylation. Moreover, mPCa cells genetically engineered to abolish *PRMT7* expression showed a reduced dissemination ability *in vitro*, *in ovo* and *in vivo*. In addition, the PRMT7 inhibitor (SGC3027) and small interfering RNA (siRNA) technology significantly suppressed the *in vitro* invasive capacity of mPCa cellular models. Furthermore, *PRMT7* was upregulated in a cohort of Spanish primary tumor samples with higher Gleason scores than in primary tumor samples with a lower Gleason scores. Therefore, our data support the role of *PRMT7* in mPCa and its potential application as a novel therapeutic target.

## RESULTS

### *In vitro* CRISPR/Cas9 screenings reveal several genes and biological processes significant for mPCa invasive process

Since the progression to metastatic cells in PCa is a complex process, we conducted two unbiased high-throughput CRISPR/Cas9 screenings to identify new genes required for metastasis onset. The GeCKO V2 Human CRISPR knockout pooled library with three pre-designed sgRNA to target almost 20,000 genes seemed the optimal choice(17). First, mPCa PC3 and DU145 cell lines with stable Cas9 expression, were generated using a lenti-Cas9 blast construct. Cas9 cells were infected with the lentiviral sgRNA library at a low Infection efficiency (MOI 0.5) to ensure single-cell infection. The library-infected pool of cells was then treated with puromycin to remove the uninfected cells.

To identify the genes that confer metastatic abilities to prostate cancer tumor cells, we conducted an invasion assay in a Matrigel-coated Boyden Chamber. DNA from cell populations that had lost their invasive abilities, which remained on top of the Matrigel membrane, and DNA from the cells that retained them, at the bottom of the membrane, were isolated separately. Subsequently, sgRNAs were amplified by PCR, sequenced, and analyzed using the MAGeCK algorithm(22), as outlined in Fig. 1A. The results of PC3 and DU145 CRISPR/Cas9 screening analyses are shown in Fig. 1B-C and Supplementary Tables 2-3.

**Figure 1.**
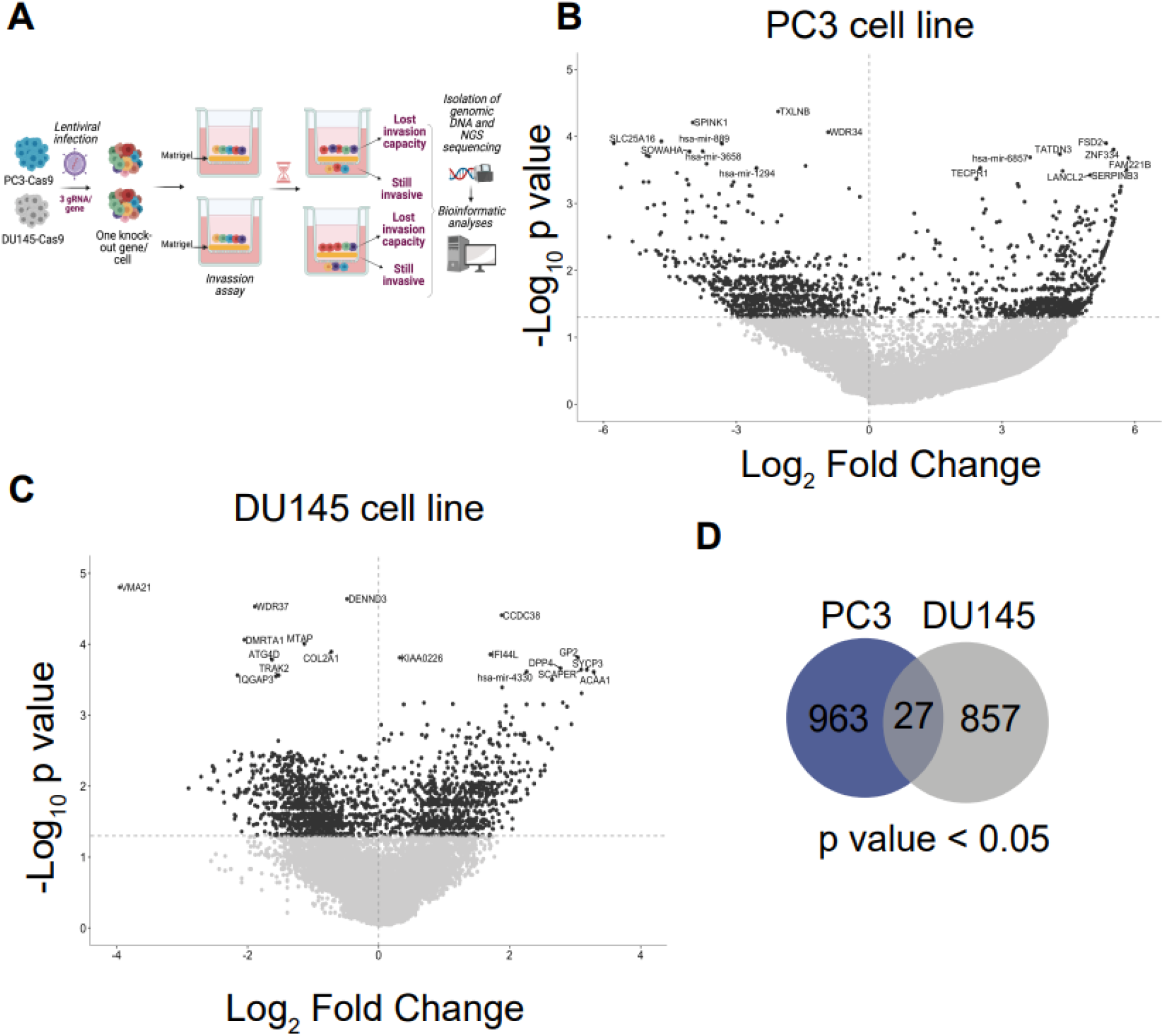
CRISPR/Cas9 screening experimental design and results. **A,** Graphical scheme of the experimental design of CRISPR/Cas9 screenings**. B,** Volcano plot showing PC3 CRISPR/Cas9 screening results. **C,** Volcano plot showing DU145 CRISPR/Cas9 screening results. **D,** Venn diagram showing the number of genes significantly associated with PCa invasive process in each line and the number of common genes between both screenings.

We were able to identify 990 genes for PC3 and 884 genes for DU145 cell line screening, which depletion significantly reduced PCa cells invasive capacities. Remarkably, 27 of them were common in both cell lines, which led us hypothesize that these genes could be more general regulators of mPCa invasion, than specific regulators of one mPCa cell line (Fig. 1D; Supplementary Table 4). Notably, we found genes such as MGAT5 that were previously associated with mPCa in a single gene study(23), proving the efficiency of CRISPR/Cas9 library technology to unbiasedly identify relevant mPCa modulators.

Furthermore, we conducted gene ontology (GO) biological pathway enrichment analyses using multiple gene functional classifications to explore whether any biological pathway was enriched in CRISPR screening results. GO enrichment analyses of PC3 screening using Metascape(24) revealed that the genes whose depletion significantly reduced invasion capacities, were mainly involved in the regulation of metabolism, the immune system, cell locomotion, and cell adhesion (Fig. 2A; Supplementary Table 5A). The same analysis in DU145 screening revealed enrichment in pathways such as the regulation of proliferation and nucleotide excision repair, the regulation of vesicle transport and the cytoskeleton reorganization or the inorganic ion homeostasis (Fig. 2B; Supplementary Table 5B).

**Figure 2.**
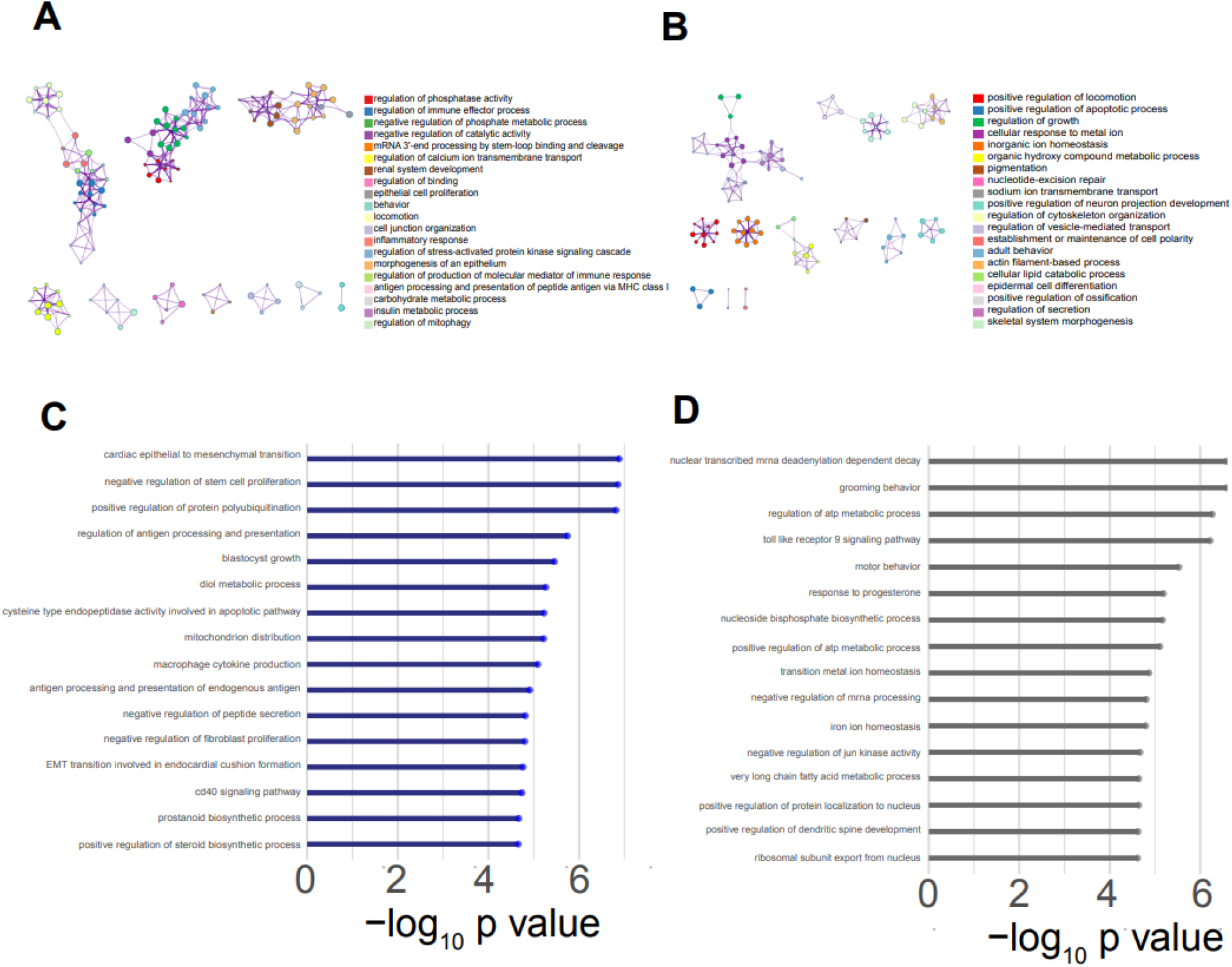
Biological pathway enrichment analyses of PC3 and DU145 screening results. A- B, Gene ontology biological pathway enrichment analysis (GO:BP) using Metascape of (24) **(A)** PC3 and **(B)** DU145 results. **C-D,** Biological pathways GSEA in **(C)** PC3 and **(D)** DU145 screening results.

Moreover, gene set enrichment analysis (GSEA) revealed an enrichment in pathways related to the epithelial-to-mesenchymal transition (EMT), the negative regulation of stem cell proliferation and the regulation of protein polyubiquitination, among others for PC3 cell screening (Fig. 2C; Supplementary Table 5C). Meanwhile for the DU145 screening, GSEA analysis showed enrichment in the mRNA nuclear processing, the ion homeostasis and negative regulation of Jun kinase, among others (Fig. 2D; Supplementary Table 5D). Overall, the biological pathways implicated in mPCa invasion varied between the two cell lines and the algorithm used, over-representation analysis or GSEA, the biological processes varied considerably. The differences observed could be due to the secondary foci from which cell line was isolated. PC3 was isolated from bone marrow metastasis(25) and DU145 from brain metastasis(26). Nevertheless, among the significantly enriched biological pathways, we found four biological pathways in common for both cell lines: the regulation of calcium ion transport into cytosol, the multicellular organism process, the DNA modification, and the DNA methylation or demethylation. Thus we hypothesized that the genes included in these biological processes are more probably general regulators of mPCa metastasis onset regardless of the organ of secondary focus formation. Additional GO enrichment analyses using Reactome and the molecular signature C5 human molecular classifications of Molecular Signatures Database (MSigDB), are shown in Supplementary Fig. 1-2; Supplementary Table 5).

### *PRMT7* ablation reduces metastatic capacities of mPCa cell lines *in vitro*

To validate our initial screening results, we selected some of our best-hit genes based on their association with other types of cancer or metastasis. We studied the *in vitro* ability of PC3 cells to invade in a Matrigel-coated Boyden chamber using fetal bovine serum (FBS) as chemoattractant, after targeting some of our best hits with siRNA. Downregulation of six of the seven candidates selected for validation led to a reduction in the invasive capacity of PC3 cells. However, this was only significant for *PRMT7*, *SYCP3* and *TECPR1* (Fig. 3A). As *PRMT7* was one of the 27 common genes found in the initial screening (Supplementary Table 4), we decided to further explore its role in mPCa. Interestingly, PRMT7 is a protein arginine methyltransferase (PRMT) that belongs to a family of other 9 proteins. It is involved in introducing ω-mono- methylation (MMA) marks in several proteins of cells. PRMT7 has been described as an important post-transcriptional regulator in breast (27) and non-small cell lung cancer metastasis(28), however, to our knowledge, its role in mPCa onset has not yet been explored. Additionally, it was one of the genes included in the biological pathway of DNA methylation and demethylation, which was significantly enriched in GSEA analyses.

**Figure 3.**
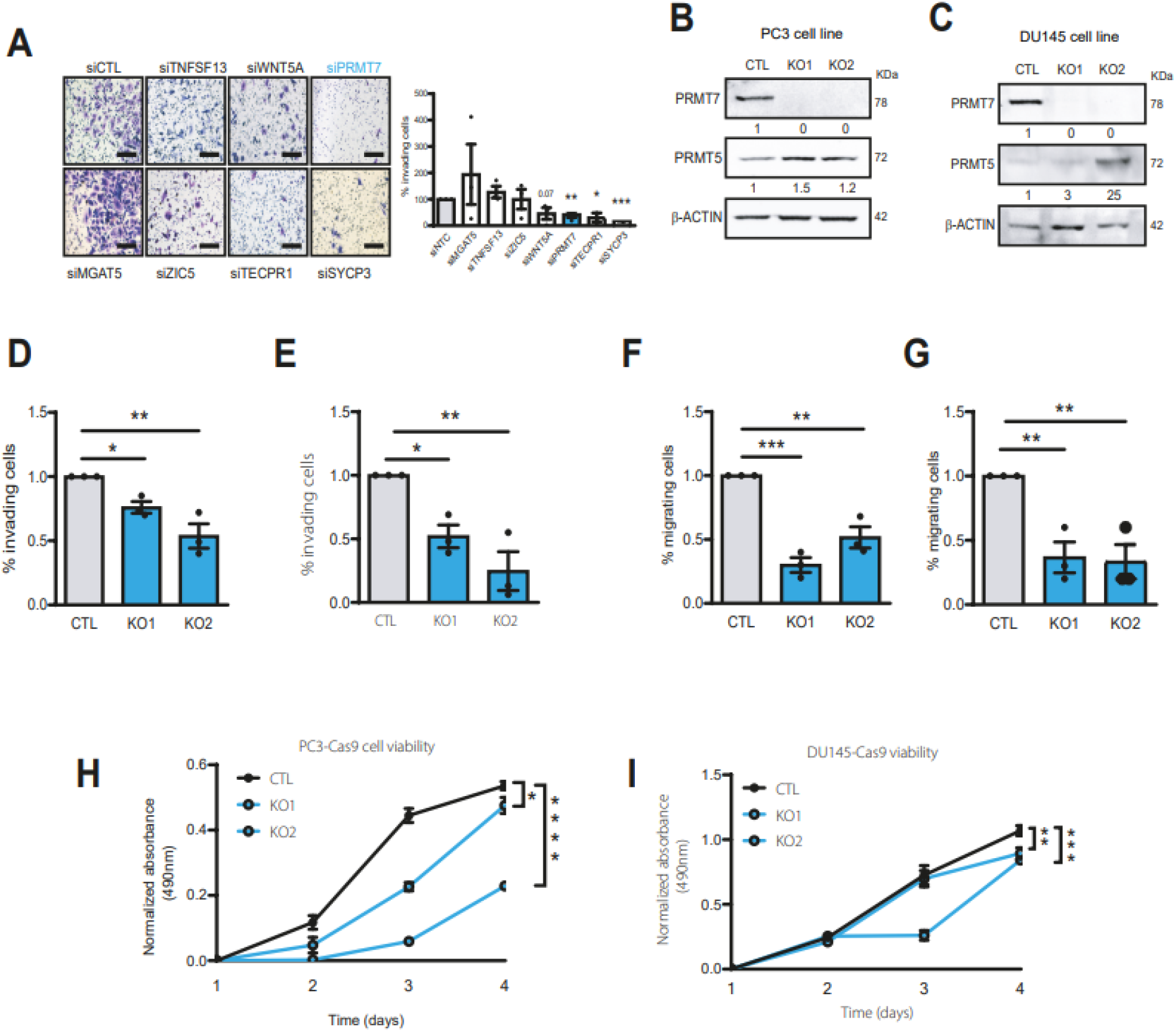
Validation of top hits by siRNA technology and further *PRMT7* role in mPCa study by CRISPR/Cas9. **A,** Invasion assay of PC3 inhibited cells using specific siRNA to target our best gene candidates versus control (siCTL) cells. **B-C,** Representative western blot of PRMT7, PRMT5 and β-actin protein levels in **(B)** PC3-Cas9 and **(C)** DU145-Cas9 cell lines. The numbers below each lane represent PRMT7/β-actin or PRMT5/β-actin respectively densitometric quantification is referred to control cells. (n=3). **D-E,** Invasion assay of *PRMT7* depleted versus control (CTL) cells of **(D)** PC3-Cas9 and **(E)** DU145-Cas9 cells (mean ± SEM of n=3 biological replicates, by unpaired Student’s t test *P < 0.05, **P < 0.01, ***P < 0.001). **F-G,** Migration assay of *PRMT7* depleted versus CTL cells of **(F)** PC3-Cas9 and **(G)** DU145-Cas9 cells (mean ± SEM of n=3 biological replicates, by unpaired Student’s t test *p < 0.05, **p < 0.01, ***p < 0.001). **H-I,** Viability assay of *PRMT7* depleted versus CTL cells of **(H)** PC3- Cas9 and **(I)** DU145-Cas9 cells (mean ± SEM of n=9 biological replicates, by TWO-way ANOVA *p < 0.05, **p < 0.01, ***p < 0.001, ****p < 0.0001).

Therefore, we decided to further analyze the role of *PRMT7* in the onset of mPCa. First, to unequivocally confirm that the effect on invasion was not due to the disruption of an off-target gene, we inhibited *PRMT7* expression by siRNA and confirmed that its silencing strongly decreased their invasive properties in both PC3 and DU145 cell lines (Supplementary Fig. 3A- D). Additionally, we analyzed the levels of MMA marks, and observed a reduction of them in the *PRMT7* depleted cells, especially in the 78 KDa band corresponding to PRMT7 auto- methylation (Supplementary Fig. 4A-B).

Next, stable PC3 and DU145 CRISPR/Cas9 *PRMT7* knock-out cell lines were generated using novel sgRNAs, different from the ones included in the GeCKO CRISPR library(17). As shown in Fig. 3B-C, western blot analyses confirmed *PRMT7* depletion in PC3 and DU145 cells without decreasing the expression of *PRMT5*, a member of the PRMT family that has been described to be able to methylate several PRMT7 targets(29).

Subsequently, we conducted invasion assays using Matrigel-coated Boyden chambers and FBS as chemoattractant for PC3 (Fig. 3D) and DU145 (Fig. 3E) cells, validating our initial screening results. We wondered whether *PRMT7* depletion could influence the ability of cells to degrade the Matrigel used in the invasion assay or it could also alter cell motility capacities. To differentiate between these effects, we conducted a migration assay in an uncoated Boyden chamber using FBS as chemoattractant, observing that *PRMT7* depleted cells had significantly lower ability to migrate than their respective controls (Fig. 3F-G) thus suggesting that *PRMT7* regulates mPCa cell migration.

Previous studies have linked *PRMT7* and proliferation(30) so we decided to study the effect of *PRMT7* depletion on the viability of mPCa cell lines viability using an MTT assay. As shown in Fig. 3H-I, a significant decrease in the viability ratio was observed in *PRMT7* depleted cells at 96h, in comparison to control cells. Hence, these results indicate that *PRMT7* silencing or depletion produces a significant reduction in PC3 and DU145 metastatic abilities, particularly in invasion, migration, and viability.

### *In vivo* model assays show reduced disseminative capacities of *PRMT7* depleted cells

*PRMT7* has been previously implicated in breast cancer metastasis(27) and non-small cell lung cancer(31) progression. Considering these studies and the *in vitro* results, we wondered whether *PRMT7* depletion plays a role in PCa distant organ colonization in animal models. To this end, we took advantage of the high metastatic potential of the PC3 cell line, and we performed two different experimental metastasis assays.

First, we conducted a chorioallantoic membrane assay (CAM)(32) as depicted in Fig. 4A. PC3- Cas9 *PRMT7* depleted and control cells were inoculated separately into the CAM of chicken embryos. Seven days later, the primary tumors generated by PC3 cells and bone marrow of chicken embryos were isolated. As shown in Fig. 4B and 4C, PC3 *PRMT7* depleted cells generated significantly smaller (Fig. 4B) and lighter (Fig. 4C) tumors than control cells. These differences in primary tumor growth *in ovo* are consistent with the decreased viability observed *in vitro*. Moreover, since PC3 cells display bone marrow tropism(33), the disseminative properties of tumor cells were assessed by qPCR, using primers against Human Alu in chicken bone marrow DNA. A lower percentage of Human Alu sequences was detected in chicken embryos inoculated with PC3 *PRMT7* depleted cells comparted to embryos inoculated with control cells, indicating that *PRMT7* depleted cells showed a lower capacity to disseminate from the primary tumor to the bone marrow of chicken embryos than control cells (Fig. 4D).

**Figure 4.**
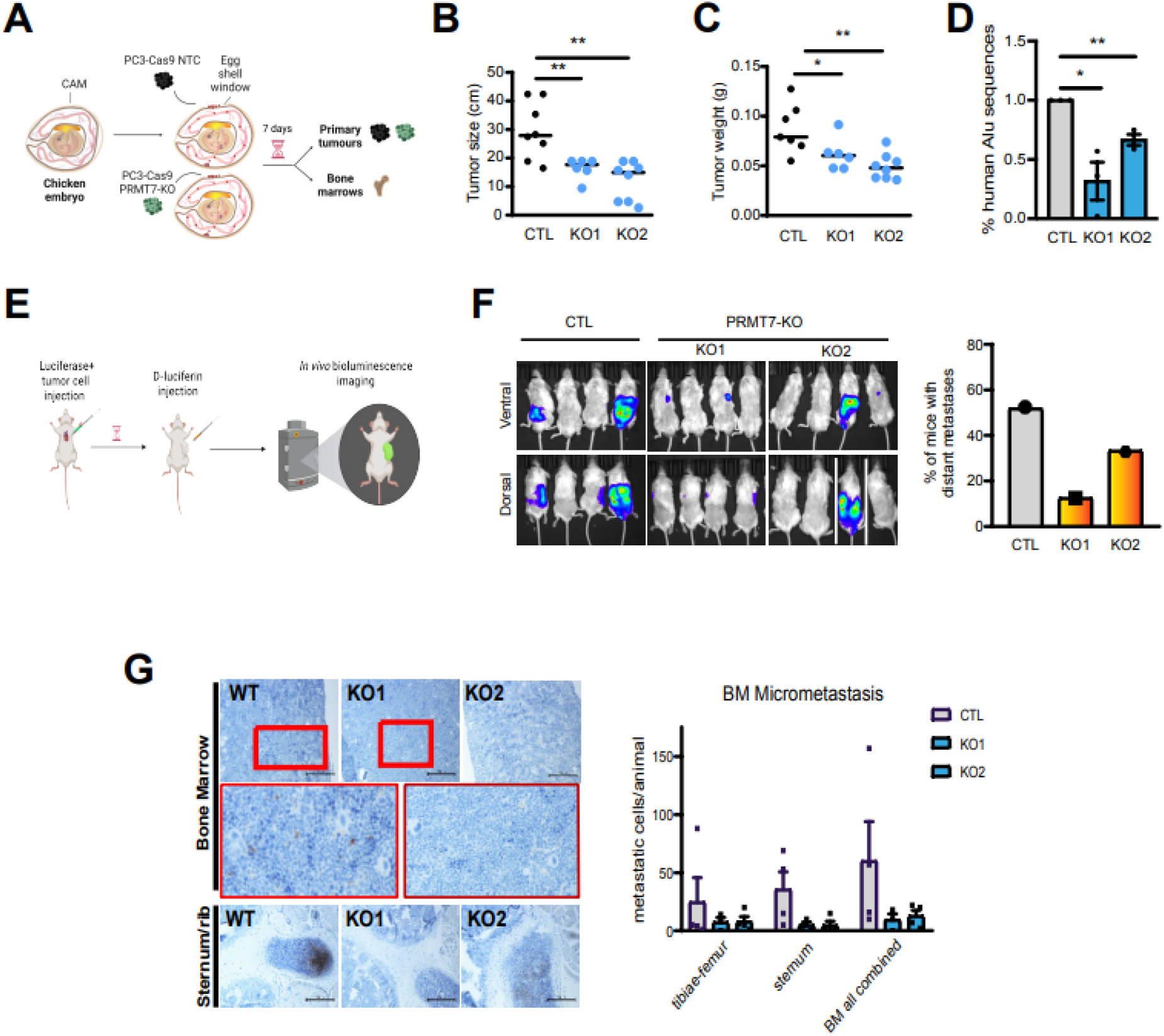
*PRMT7* depletion reduces proliferative and disseminative abilities of PCa cells *in ovo* and *in vivo.* **A,** Graphical scheme of *in ovo* studies experimental design. **B-C,** Graphs representing primary tumor **(B)** size and **(C)** weight (mean ± SEM of n=6-8 inoculated chicken embryos, by ordinary ONE-way ANOVA *p < 0.05, **p < 0.01, ***p < 0.001). **D,** Graph representing percentage of human Alu sequences amplified by qPCR in chicken bone marrow (mean ± SEM of n=3, Student’s t test). **E,** Graphical scheme of *in vivo* dissemination assay experimental design**. F,** Left panel shows representative pictures of metastases generated by CTL, *PRMT7*-KO cells. Right panel shows the percentage of mice with distant metastasis (n=12). **G**, Left panel shows representative immunohistochemistry images of GFP-positive cells in the muse bones. Right panels show the percentage of disseminated tumor cells (DTC) in mouse bone marrow (mean ± SEM of n=9 animals, by Student’s t test).

Previous studies have shown that upon direct inoculation of PC3 cells in the blood circulation of immunodeficient mice (NOD-SCID), they were able to generate metastatic foci in distant tissues including the femur, tibia, jaws, and ribs(33). Hence, we inoculated control and *PRMT7* depleted PC3 cells engineered to express a luciferase-GFP DNA construct using a lentiviral approach. They were inoculated into the left cardiac ventricle of athymic nude mice, as previously described(34). Mice were analyzed using an IVIS system and sacrificed after 28 days, and their tissues (including femur, tibiae, and sternum) were harvested, fixed, and analyzed for the presence of micrometastatic foci, as depicted in Fig 4E. Distant metastasis was observed in 50% of mice inoculated with control cells compared to 12% or 33% of mice inoculated with *PRMT7*- KO1 and *PRMT7*-KO2 depleted PC3 cells, respectively. A representative image is shown in Fig. 4F. Unexpectedly, most of the detected macrometastases displayed visceral localization, whereas no bone macrometastases were detected (Supplementary Fig. 5). To analyze the presence of micrometastases of PC3 cells in the bone marrow immunohistochemistry assays of bone marrow from the tibia, femur, and sternum were performed. Quantification of disseminated tumor cells by GFP staining revealed that *PRMT7* depletion in PC3 cells severely reduced the formation of bone marrow micrometastases compared to that in PC3 control cells (Fig. 4G). These results highlight the relevance of PRMT7 in controlling biological processes that are directly implicated in PCa metastasis onset both *in vitro* and *in vivo*.

### Expression of relevant cell adhesion molecules is regulated by *PRMT7*

To elucidate the biological mechanisms controlled by PRMT7 that facilitate distant organ colonization of primary tumor PCa cells, we conducted differential expression analysis of PC3 *PRMT7* depleted versus control cells. Out of the total genes analyzed, the expression of 781 genes was significantly altered (Fig. 5A; Supplementary Table 6). Among the most differentially expressed genes, we found *PNPLA4*, *SFMBT2,* and several genes related to cell adhesion such as *TUBAC3* and *CHST15*. *TUBAC3* encodes a protein implicated in microtubule formation, whereas *CHST15* encodes a glycosaminoglycan that binds to proteins to generate proteoglycans, which are important components of the extracellular matrix.

**Figure 5.**
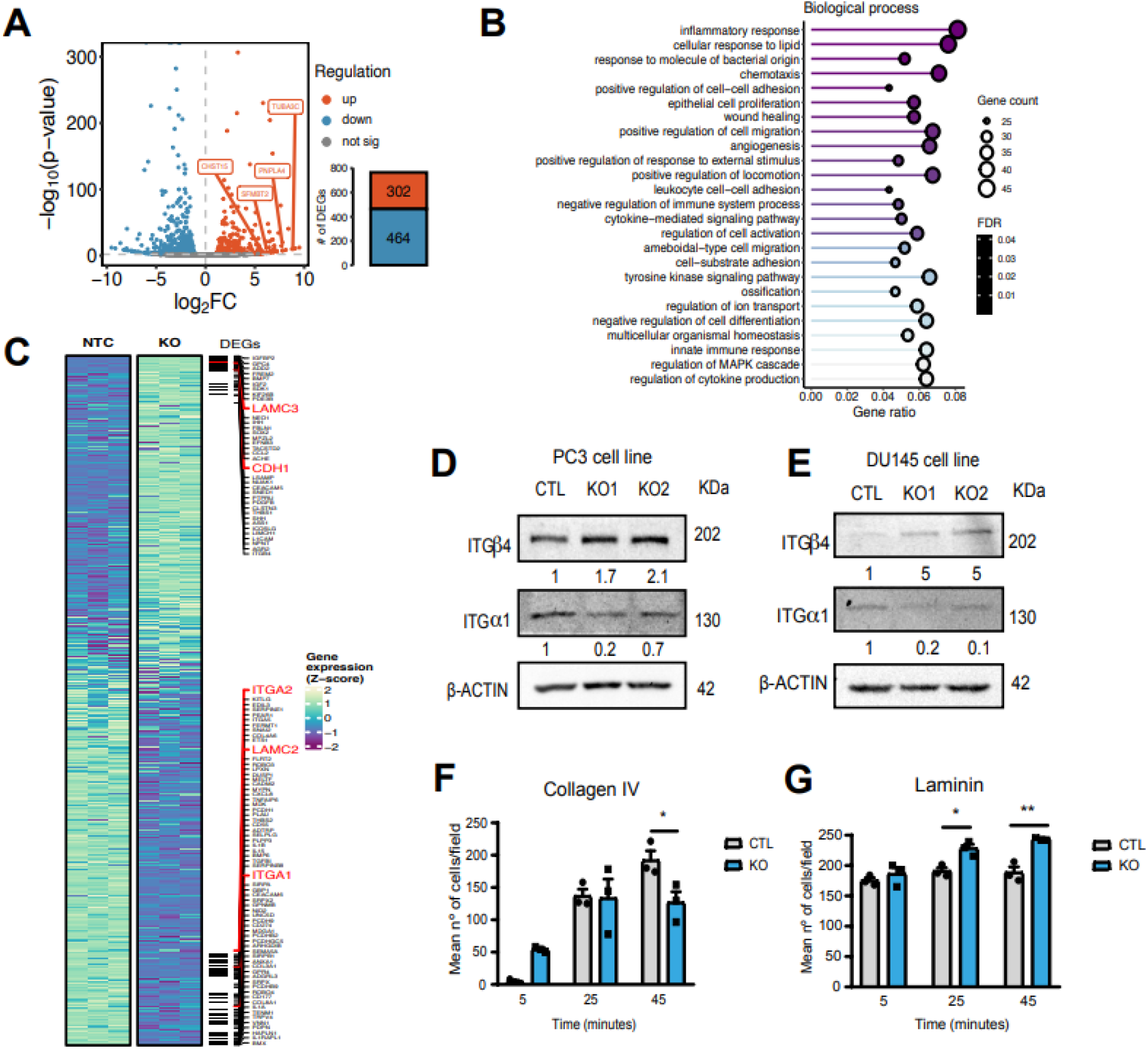
*PRMT7* depletion leads to an adhesion molecule switch in PC3 cells. **A,** Volcano plot showing differentially expressed genes (DEGs) (adjusted p-value < 0.05, |logFC| > 1) from the differential gene expression analysis of PC3 control versus *PRMT7* depleted cells results, and a barplot representing the number of genes that significantly changed their expression, accounting for the number of genes that were upregulated (red) and downregulated (blue) in *PRMT7* depleted cells. **B,** Over-representation analysis of biological process GO terms in DEGs (adjusted p-value < 0.05, |logFC| > 1) of *PRMT7* depleted cells compared to control cells, representing top 25 terms with highest gene ratio with an adjusted p-value cutoff of 0.05 and redundant GO terms were removed. **C,** Heatmap showing the gene expression (Z-score) of the genes found in parental GO term cell adhesion sorted by log_2_ fold change. Horizontal lines denote DEG positions, and those of interest are highlighted in red. **D-E,** Western blot analysis of ITGα1 and ITGβ4 protein levels in **(D)** PC3-Cas9 and **(E)** DU145-Cas9 cells. The numbers below each lane represent ITGα1/β-actin or ITGβ4/β-actin densitometric quantification referred to control cells, respectively. **F-G,** Graph representing mean number of cells per field adhered to **(F)** collagen IV or **(G)** laminin (mean ± SEM of n=3, by unpaired Student’s t test *P<0.05, **P>0.01).

Consecutively, a GO over-representation analysis was performed, as shown in Fig. 5B, which revealed that the cell-substrate adhesion term was over-represented. Hence, we explored the overall expression of genes annotated to the cell adhesion parental ontology term (GO:0007155) (Fig. 5C). Surprisingly, we observed that some genes, such as *CDH1* or *LAMC3*, were upregulated in *PRMT7* depleted cells, while others such as *ITGA1, ITGA2* and *LAMC2,* were downregulated.

To validate our RNA-seq results, we randomly selected some of the genes included in the cell adhesion group and analyzed their expression in independent PC3 *PRMT7* depleted and control samples using RT-qPCR, as shown in Supplementary Fig. 6. Cell adhesion has been described as a crucial biological process that mediates cell motility and colonization of distant organs, with integrin receptors being especially relevant(35). Moreover, the overexpression of some integrins has been associated with poor prognosis in several cancers, and depending on the type of integrin expressed on their membranes, cells can adhere better to specific types of extracellular matrixes(36). Thus, we analyzed the protein levels of ITGα1 and ITGβ4 that, mediate binding to collagen IV or laminin matrices, respectively.

Western blot analyses revealed that ITGα1 protein levels were significantly reduced in *PRMT7* depleted cells, while ITGβ4 protein levels were significantly increased in comparison to control cells, in both, PC3 (Fig. 5D) and DU145 (Fig. 5E) cell lines. Interestingly, overexpression of exogenous *PRMT7*(37) reduced ITGβ4 levels in both control and *PRMT7* depleted cells, which is in agreement with previous results (Supplementary Fig. 6). Hence, cell adhesion to collagen IV (Fig. 5F) and laminin (Fig. 5G) was evaluated by observing a change in selective adhesion to these matrices. Overall, PC3 cells, both *PRMT7* depleted and control, had a greater ability to adhere to laminin than to collagen IV. However, we observed that *PRMT7* depleted cells had a higher adhesion capacity than the control cells. In addition, at longer time points, control cells preferentially adhered to collagen IV, while *PRMT7* depleted cells maintained better adhesion to laminin (Fig. 5F-G). These results indicate that overexpression of *PRMT7* in primary tumor cells could produce a switch in the expression of adhesion molecules, leading to a reduction in cell adhesion while increasing cell motility and, promoting dissemination to distant organs.

### Arginine methyltransferase 7 regulates the activity multiple transcription factors

PRMT7 is present in the nucleus and cytoplasm, introducing post-transcriptional modifications to several proteins(29). Hence, we investigated how PRMT7 modulates the expression of cell adhesion genes identified in the transcriptomic analysis. Accordingly, we investigated the transcription factors (TFs) that might regulate these transcriptome differences using two alternative approaches. First, we performed an enrichment analysis of TF binding sites within the promoters of differentially expressed genes in PC3 cells, which revealed several TF related to Polycomb complex-2, such as EZH2 (Fig. 6A). This histone methyltransferase has already been associated with PCa progression(12).

**Figure 6.**
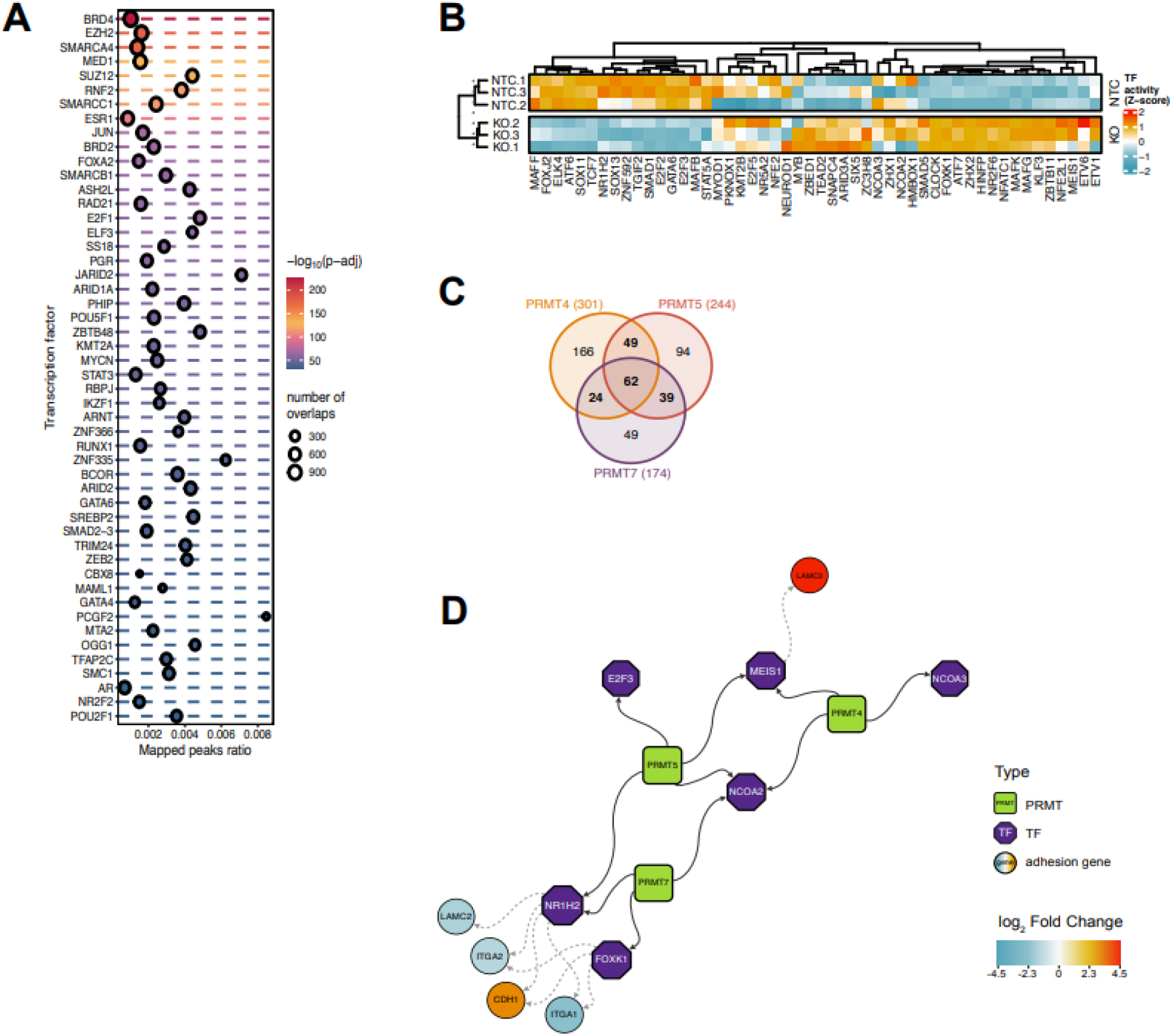
PRMT7 drives cell adhesion molecule reprogramming through several TF methylation. **A,** Top 50 most significant enriched TFs in the promoter regions of differentially expressed genes (DEGs). **B,** Heatmap showing TF activity estimated from TF regulons expression, representing the top 50 most variable TFs across CTL and *PRMT7* depleted cell samples. **C,** Venn diagram of the target proteins of PRMT7, PRMT4 and PRMT5 described in Li *et al*^36^. **D,** Regulatory network of transcription factors (purple hexagons) that are methylated by PRMT7, PRMT4 and PRMT5 (green squares), and can bind to cell adhesion gene promoters (circles).

Alternatively, we also inferred TF activities based on the expression values of their target genes for each sample and selected the top 50 TFs, with the highest variation across samples independent of their experimental treatment. Our results revealed that the most variable TFs showed high concordance with changes in TF activity between *PRMT7* depleted and control cells (Fig. 6B). Surprisingly, we identified 7 of top 50 TFs with arginine methylation sites from dbPTM database, and intriguingly 6 of them presented a robust increased activity on PRMT7 depleted samples (Fig. 6B). Curiously, we found that 4 of those 7 TFs (FoxK1, NR1H2, NCOA2 and NCOA3) were PRMT7s targets as described in the recently published PRMT4, PRMT5 and PRMT7 methylomes(38). In Fig. 6C is depicted a Venn diagram showing the proteins susceptible of being methylated by the three, two or one of the PRMTs. Therefore, we focused our analysis on the TFs controlling the expression of cell adhesion-related genes susceptible to PRMT7 methylation. FoxK1, NR1H2, and NCOA2/3 are the only TF susceptible to specific methylation by PRMT7(38), and are able to bind to the promoter of cell adhesion genes, including the analyzed *CDH1* and *ITGA1/2*, where they could promote or inhibit their expression, as summarized in Fig. 6D and Supplementary Fig. 7. Altogether these results suggest that *PRMT7* depletion alters the activity of relevant transcription factors of cell adhesion genes such as *FoxK1*, *NR1H2,* and *NCOA2/3,* leading to reprogramming of the type of cell adhesion molecules expressed on the cell surface.

### Potential use of *PRMT7* as a new therapeutic target or biomarker

To further explore the role of PRMT7 in PCa metastasis, we analyzed *PRMT7* gene expression levels in a cohort of Spanish FFPE-embedded PCa primary tumor samples recruited from Hospital Clínico San Carlos (Madrid, Spain). From the total of samples analyzed (n=13), patients without *de novo* PCa metastasis harboring a 6 or lower Gleason score (GS) were labbeled as the “Low” group meanwhile patients diagnosed with *de novo* PCa metastasis and with 7 or higher GS were classified as “High” group. Table 1 summarizes of the patient characteristics included in the study, and Fig. 7A shows representative H&E staining pictures of primary tumor samples obtained from PCa patient diagnosed with low and high risk PCa tumors. As shown in Fig. 7B, samples obtained from patients diagnosed with *de novo* metastatic disease and high GS showed higher levels of *PRMT7* expression than non-metastatic and low GS samples. These results suggest that *PRMT7* may be a potential biomarker of PCa malignancy.

**Figure 7.**
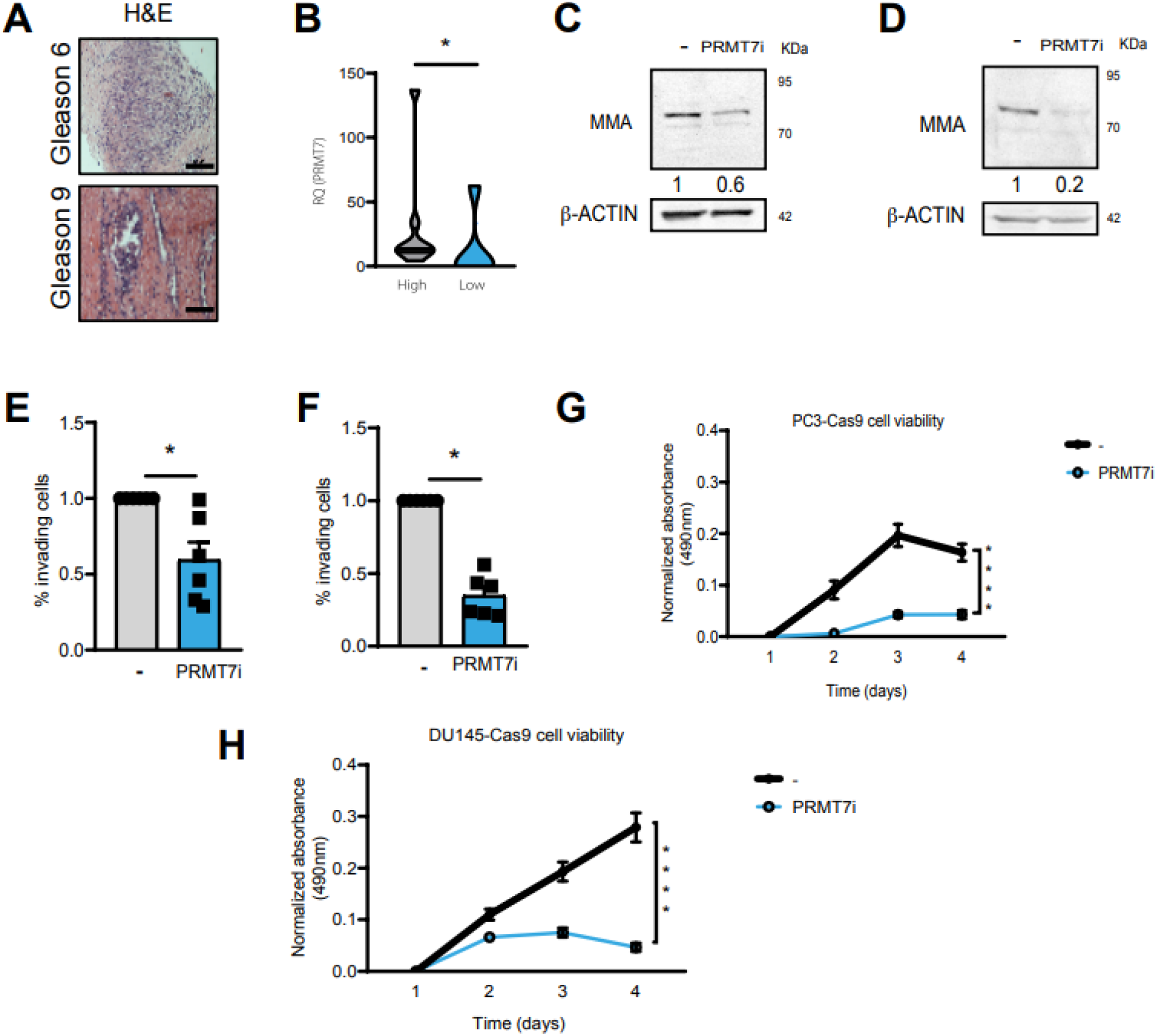
PRMT7 clinical approach. **A** Hematoxylin and eosin staining of PCa primary tumor slides showing the aggressiveness of tumors measured by Gleason score *PRMT7* (scale bars: 50 μM). **B,** *PRMT7* gene expression levels in PCa primary tumor samples with higher (n=8) versus lower (n=5) Gleason score samples (mean ± SEM of n=13 primary tumor samples, by unpaired Mann-Whitney test *p < 0.05, **p < 0.01, ***p < 0.001). **C, D,** Representative western blot of arginine monomethylation (MMA) levels in **(C)** PC3 and **(D)** DU145 cells treated with 10μM of SGC3027 (PRMT7 inhibitor). The numbers below each lane represent MMA/β-actin densitometric quantification referred to control cells. **E-F,** Invasion assay of PRMT7 inhibited versus vehicle cells of **(E)** PC3 and **(F)** DU145 cells (mean ± SEM of n=6 biological replicates, by unpaired Student’s t test *p < 0.05, **p < 0.01, ***p < 0.001). **G-H,** Cell viability assay of PRMT7 inhibited versus vehicle cells of **(G)** PC3 and **(H)** DU145 cell lines (mean ± SEM of n=9 biological replicates, by TWO-way ANOVA *p < 0.05, **p < 0.01, ***p < 0.001, ****p < 0.0001).

**Table 1.** Summary of PCa patient clinical data. In the table are summarized the mean age, Gleason score or OMS group and the percentage of tumor of FFPE-embedded samples from Hospital Clínico San Carlos.

| Characteristic | Low Gleason Score cohort (n=5) | High Gleason score cohort (n=8) | Both groups combined (n=13) |
| --- | --- | --- | --- |
| Age at time of treatment, mean (min-max) | 67 (53-72) | 69 (60-85) | 68 (53-85) |
| Gleason score, mean (min-max) | 6 (6-6) | 8 (7-10) | 7 (6-10) |
| OMS group | 1 (1-1) | 4 (2-5) | 3 (1-5) |
| Percentage of tumor in the sample (%), mean (min-max) | 7% (0%-20%) | 50% (20%-90%) | 33% (0%-90%) |

Our *in vitro*, *in vivo* and *PRMT7* expression analysis in PCa patients taken together with the fact that some inhibitors of PRMT family are already in clinical trials(39), we decided to analyze the effect of PRMT7 inhibitors on the metastatic abilities of our mPCa cell lines. First, we checked the state of mono-methylation (MMA) levels in cells after treatment with the SGC3027 inhibitor (10 μM), as shown in western blots of Fig. 7C for PC3 and Fig. 7D for DU145 cells. We observed a reduction in MMA levels in the cells treated with the PRMT7 inhibitor (PRMT7i) in comparison to cells treated with vehicle, which suggests that PRMT7 activity was reduced when the cells were treated with the inhibitor. The next step was to check whether pharmacological inhibition was sufficient to reduce the invasive capacity of mPCa cells using FBS as chemoattractant. As shown in Fig. 7E-F, pharmacological inhibition of PRMT7 in PC3 (Fig. 7E) and DU145 (Fig. 7F) cells led to a significant decrease in *in vitro* cell invasion compared to that in cells treated with the vehicle. Moreover, we checked the viability of cells treated with the PRMT7 inhibitor versus those treated with the vehicle. A reduction in proliferation and/or viability, mainly at 96h was observed in PC3 (Fig. 7G) and DU145 (Fig. 7H) cells treated with the inhibitor. Altogether, these results suggest that pharmacological PRMT7 inhibition could be an effective approach to decrease the metastatic capacity of PCa cells *in vitro* and open a field of research for the future at an *in vivo* preclinical level.

## DISCUSSION

Metastasis is a complex phenomenon involving many biochemical processes that are largely unknown(8). Although there are several effective therapies for the initial stages of PCa, metastatic castration-resistant prostate cancer has not a cure yet. The high incidence of this neoplasia highlights the need to identify new targets for use as prognostic biomarkers and/or therapeutic targets. To our knowledge, our report presents the first two independent large-scale screenings specifically designed to uncover PCa invasion gene drivers using CRISPR/Cas9 based technology. Bioinformatic analysis, allowed us to identify many novel genes and biological pathways essential for PCa cell invasion.

Our extended *in vitro* and *in vivo* characterization of PRMT7 revealed that this gene is a critical regulator of PCa tumor progression, being relevant not only for cell viability of the cells but also for regulating the invasive and migratory ability of cells, which are key to metastasis development. According to our results, PRMT7 controls PCa cell viability, both *in vitro and in vivo.* However, the reduction in viability upon *PRMT7* depletion would not explain the decrease in PC3 and DU145 motility, as differences in migration and invasion were measured at 24h, while viability differences were observed at longer time frames. Moreover, we validated these results using PRMT7 pharmacological inhibitors, which could be used to prevent the appearance of mPCa.

PRMT7 is the only member of the arginine methyltransferase family belonging to subgroup III and is involved in the preferential introduction of MMA(29). In the context of breast cancer, PRMT7 has been shown to methylate the E-cadherin proximal promoter, inducing EMT transition(27); to upregulate the expression of matrix metalloproteinase-9 (MMP9)(40); and to methylate SHANK2, activating endosomal FAK signaling(41) which all together lead to the progression of the disease. In non-small cell lung cancer, PRMT7 contributes to metastasis appearance through interaction with HSPA5 and EEF2(28). In addition, it has been described that in renal cell carcinoma, PRMT7 can methylate β-catenin promoting its stabilization and the induction of c-myc expression, which is an important regulator of cell proliferation and tumor progression(30). Moreover, PRMT7 has been described to methylate the Wnt signaling molecule Dishevelled 3(42), the transcription factor C/EBP-β(43), and the RNA splicing factor hnRNPA1(38), suggesting that it, directly or indirectly, could be implicated in several cellular pathways, both in cellular homeostasis and disease. Therefore, due to the high number of biological processes susceptible to PRMT7 methylation, we decided to conduct a differential transcriptomic analysis in our mPCa cellular models (PC3 *PRMT7* depleted versus control cells) to elucidate the most important processes regulated by PRMT7, which are crucial to mPCa onset. As expected, the results showed differences in *CDH1* expression. However, we did not detect significant changes in the expression of other genes, such as *MMP9*, *HSPA5*, *EEF2, CTNNB1*, nor *MYC*. Remarkably, we observed and enrichment of several cell adhesion biological processes.

Cell adhesion is a widely known mechanism that contributes to tumor cell migration and invasion. In fact, changes in the expression of some integrin and laminin receptors, which are well-characterized cell adhesion molecules, are already considered malignant biomarkers. For example, the alpha 1 subunit of integrin receptors (*ITGA1)* is a pre-malignant biomarker that promotes therapy resistance and metastatic potential in pancreatic cancer(44) whereas laminin subunit gamma 2 (*LAMC2*), has previously been identified as a specific marker for metastatic abilities in lung adenocarcinoma(45). Therefore, we focused our functional validation in demonstrating that *PRMT7* depletion leads to a switch in the expression of some integrins, laminins, and cadherins, such as *ITGA1*, *ITGB4*, *LAMC2*, *LAMC3* and *CDH1,* which consequently alters the adherence of mPCa cells to different types of extracellular matrices, such as collagen IV and laminin, and that at least ITGβ4 and ITGα1 changes are reversed upon exogenous *PRMT7* expression. Bone marrow is an organ rich in collagen IV, therefore we suggest that cells with higher levels of PRMT7 could adhere and survive better in this secondari focus, where 70% of PCa metastasis are found.

In this study, TF activity analysis uncovered a new role for PRMT7 as a regulator of several TFs. These results, together with the previously published PRMT7 methylome(38), suggest that either methylation of FoxK1, NR1H2, or NCOA2/3 could regulate their transcriptional activity in mPCa cells. Remarkably, FoxK1, a TF susceptible to methylation by PRMT7 on R161 and R191 residues, is a known regulator of the expression of several adhesion molecules and has previously been involved in the acquisition of mPCa abilities(46),(47). In addition, NR1H2 is susceptible of being methylated on R126 and is known to regulate the expression of LAMC2 and CDH157(48). Arginine methylation stabilizes E2F1(49); therefore we speculate that it may have a similar effect on FoxK1, NR1H2, or NCOA2/3 allowing primary PCa cells to gain migratory and invasive abilities and enhance distant organ colonization. Regulation of cellular adhesion molecules via PRMT7-FoxK1/NR1H2/NCOA2/3 may explain the acquisition of migratory abilities shown in PCa cellular model studies.

Regarding the use of PRMT7 as prognostic biomarker or therapeutic target, although further validation analyses with larger cohorts of samples are needed, results are promising. In fact, some studies have already highlighted PRMTs as an emergent group of proteins involved in cancer and metastasis(50). Previous works have described other members of the family, especially PRMT5, as important cancer regulator(51). Indeed, PRMT1 and PRMT5 inhibitors are currently under Phase I clinical trials (NCT04676516, NCT03573310, NCT05094336, NCT03854227, NCT04089449, NCT03886831, NCT05275478, NCT04794699, and NCT05245500). However, the role of PRMT7 in mPCa has not been explored. While previous studies have linked its overexpression to breast cancer(52), non-small cell lung cancer(28) and renal carcinoma progression(30), to our knowledge, this is the first study to unravel the association between PRMT7 and PCa progression.

In summary, we conducted two large-scale CRISPR/Cas9 library screenings to identify the most relevant drivers of PCa cell invasion and validated some of the most robust gene candidates, with being *PRMT7* the best candidate. Here, we show that its inhibition by siRNA or pharmacological approaches, or its depletion by CRISPR/Cas9, significantly reduces the invasion, migration, and viability of mPCa cell lines *in vitro* and *in vivo.* Furthermore, we found in a cohort of Spanish PCa samples that it is upregulated in primary tumors with a high Gleason score compared to less aggressive tumors. Moreover, our transcriptomic analysis indicated that this arginine methyltransferase could do so through the methylation of several TFs, such as FoxK1 or NR1H2, which could consequently alter the expression of certain laminins, integrins and cadherin molecules. Taken together, these results highlight *PRMT7* as a promising biomarker candidate for poor prognosis and a potential new therapeutic target for the treatment of PCa metastasis.

## MATERIALS AND METHODS

### Cell culture and reagents

PC3 (CRL-1435) and DU145 (HTB-81) cell lines were obtained from ATCC and cultured in RPMI/HAM’S F12 and RPMI medium supplemented with 10% FBS, respectively. HEK293T (CRL-3216) cells obtained from ATCC, were cultured in DMEM medium supplemented with 10% FBS and were used for lentiviral production.

### CRISPR/Cas9 library preparation

The lentiCRISPR v2 GeCKO Human library (Addgene plasmid #52961) was amplified following Sanjana et al., protocol(17). For viral production, see Supplementary Material Methods. PC3-Cas9 and DU145-Cas9 cells were transduced with lentivirus containing pre-designed sgRNAs in the presence of 8 μg/mL polybrene. 48h later, puromycin selection (10 μg/mL) was performed for 5 days. Next, an invasion assay in Matrigel-coated Boyden chambers was conducted separately for both cell lines.

### MAGeCK screening analysis

DNA from the cell population in the upper chamber of the Matrigel-coated Boyden Chamber membrane and the cell population at the bottom of the membrane, was isolated separately. Specific primers designed to amplify all sgRNAs included in the lentiCRISPR v2 GeCKO Human library (Supplementary Table 1) were used to amplify the sequences that were included in the DNA of the cells through lentiviral infection. Subsequently, next generation sequencing (NGS) and analysis of raw data were conducted at the Bioinformatics for Genomics and Proteomics Unit at the CNB-CSIC facility (Madrid, Spain) using the MAGeCK algorithm(22). To proceed with further analysis, we discarded the genes in which only one sgRNA was detected by NGS to improve the reliability of the screening results.

### Gene silencing by small interfering RNA

Pre-designed siRNAs were obtained from Dharmacon (Supplementary Table 1). 2.5 × 10^5^ cells were initially transfected with 50 nM siRNA using Lipofectamine RNAiMAX following the manufacturer’s protocol (Invitrogen). After 48h of incubation, cells were used for invasion or immunobloting assays.

### PRMT7 CRISPR/Cas9 knock-out production

Lenti-sg*PRMT7* and lenti-sgControl were performed as previously described(14), using the specific guide *RNA* for *PRMT7* (forward 5’-*CACCG*AAGGCCTTGGTTCTCGACAT-3’ and reverse 5’- *AAA*CATGTCGAGAACCAAGGCCTTC-3’) or for the control (sgCTL) (forward 5’- *CACCG* CGGCTGAGGCACCTGGTTTA-3’ and reverse 5’- *AAA*CTAAACCAGGTGCCTCAGCCG-3’). PC3-Cas9 and DU145-Cas9 previously engineered in the laboratory, were infected with *sgPRMT7* or sgCTL in the presence of polybrene (8 μg/mL). After 48 days, puromycin selection (10 μg/mL) was performed for 5 days. Serial dilutions were performed in 96 multi-well plates to obtain single-cell clones. *PRMT7* knock-out was confirmed by western blot using anti-PRMT7 antibody and by Sanger sequencing (data not shown).

### Protein extraction and immunoblotting

Cells were lysed and western blot protocol was carried out as previously described(53). The primary antibodies used were β-Actin (sc-47778, Santa Cruz Biotechnology), PRMT7 (#14762S, CST), mono-methyl arginine (Me-R4-100, CST) (#8015S, CST), PRMT5 (sc-376937, Santa Cruz Biotechnology), ITGα1 (sc-271034, Santa Cruz Biotechnology), and ITGβ4 (sc-9090, Santa Cruz Biotechnology). The secondary antibodies used were anti-mouse IgG (NA931, Cytiva) or anti-rabbit IgG (NA934, Cytiva).

### Invasion, migration and cell adhesion assays

Medium supplemented with 5% of FBS was placed in the lower chamber and used as chemoattractant in both experiments. Invasion was assessed in Matrigel (30μg; 45356231, Corning) coated transwells. 7.5 x 10^4^ cells were seeded in the upper chamber in serum-free medium. For migration assessment, 2.5 x 10^4^ cells were seeded in serum-free medium in the upper chamber of a 48-Well Micro Chemotaxis Chamber (NeuroProbe, AP48). After 24h, invading cells were fixed with 4% paraformaldehyde and stained with 0.2% crystal violet or fixed and stained using the Kwik Diff kit (Epredia), respectively. Images were taken with an Eclipse TE300 Nikon microscope and quantified using ImageJ software. Invasion experiments to evaluate the effect of SGC3027 PRMT7 inhibitor (SML2343, Sigma-Aldrich) were carried out pre-treating cells with 10μM of inhibitor overnight. Cell adhesion was performed as previously described(14), using pre-covered wells with laminin or collagen IV (5μg/cm^2^).

### MTT assay

Cell viability was assessed using the CellTiter 96 Aqueous One Solution Cell Proliferation Assay Kit (Promega) following the manufacturer’s protocol, as previously described(14). To assess the relevance of the SGC3027 pharmacological inhibitor in PCa cells, cells were cultured in 10% FBS media with 10 μM (SML2343, Sigma-Aldrich).

### Metastasis assays in chicken embryos

Experiments were performed as described previously(32). Briefly, 1x10^6^ PC3-Cas9 sgCTL or PC3-Cas9 *PRMT7* KO1 and KO2 were inoculated into the chorioallantoic membranes (CAMs) of embryonic day 10 (E10) chicken embryos. On day 7 (E17) after inoculation, primary tumors and bone marrow of the embryos were isolated. Primary tumors were sized and weighted, and DNA was extracted from the bone marrow using XNAT2-1KT kit (Sigma Aldrich). The presence of human Alu sequences in the DNA of chicken bone marrow was analyzed by qPCR as described previously. Primer sequences are listed in Supplementary Table 1.

### Tumor growth and bone metastasis in nude mice

Mice studies were carried out in strict compliance with the European Community Council Directive (2010/63/EU) and following guidelines for animal research from the Complutense University of Madrid (UCM) Ethical Committee, approved by the Community of Madrid (Spain) with reference (PROEX 127.0/21). Bone metastasis experiments were performed as previously described(33). Briefly, PC3 cell lines, both control and PRMT7 knock-out, were infected with GFP-Luciferase Lentivirus and sorted. We injected 10^5^ cells into the left cardiac ventricle of male NOD SCID J mice (Charles River Laboratories). The appearance of mouse metastasis was monitored by IVIS (IVIS Lumina III, Perkin Elmer) after injecting the animals with luciferine (150 mg/kg, Biotherma).

For routine histological analysis, bones were fixed in 10% buffered formalin (Sigma), decalcified in 0.5 mol/L EDTA for 7 days, incubated in 30% sucrose, and embedded in paraffin. 2μm paraffin sections were subjected to immunohistochemical analysis using an antibody against GFP (#2956, CST). The tissue slides were scanned using an AxioScan Z1 scanner (Zeiss). Digital images were analyzed, and the number of micrometastases was determined in one or two slides from four randomly chosen mice per group. Microscopy and software calibration for size measurements were conducted using a TS-M2 stage micrometer (Oplenic Optronics).

### RNA isolation and sequencing

Total RNA from three replicates of *PRMT7*-KO2 and control PC3 cells was extracted using the NucleoSpin RNA kit (Macherey-Nagel). mRNA quality control checks and NGS for RNA-seq analysis were performed at the NIMgenetics facility (Madrid, Spain). To validate the RNA-seq results, reverse transcription (RT) was performed using Superscript IV Reverse Transcriptase (ThermoFisher). Triplicate samples with their corresponding controls were assessed by qPCR, performed at the UCM Genetic and Genomic Facility. Gene fold changes were determined using the 2-^ΔΔ^Ct algorithm. All the primer sequences are listed in Supplementary Table 1. GAPDH was used as the reference gene to normalize gene expression.

### RNA-Seq data processing and analysis

RNA-Seq data in FASTQ format were mapped against the reference human genome GRCh38.p13 using STAR v2.7.9a(54) and GENCODE V38 annotation as a gene model that included a total of 60,649 annotated genes. Gene-level abundances were counted using HTSeq(55), software that is internally available in STAR. Downstream analyses were performed using R v4.1.2(56). Lowly expressed genes were filtered out when accounting for less than 15 read counts across all samples.

### Differential gene expression analysis

Differential expression analysis was carried out using DESeq2 v1.34.0(55) with the design formula ∼ Condition, a factor with two different levels (i.e., sg*PRMT7*-KO, sgCTL). Genes were considered differentially expressed (DEG) using a 5% FDR and an absolute log2 Fold Change > 1 as thresholds.

### Gene Ontology overrepresentation analyses

Gene ontology (GO) overrepresentation analysis of biological process terms was performed using clusterProfiler v4.4.2(57) and org.Hs.eg.db v3.14.0(58). Redundant GO terms were excluded for graphical representation using the rrvgo v1.6.0(59), that is based in semantic similarity and adjusted p-value.

### TF enrichments

Transcription factor (TF) enrichment was calculated within the promoters of DEGs using ReMapEnrich v0.99.0(60) which computes overlappings TF binding sites (established in non-redundant peaks ReMap2022 hg38 catalog(60)) and promoter regions. Gene promoters were established at 2500 upstream and 500 downstream base pairs of the TSS using the GenomicRanges v1.46.1(61).

### TF activity based on gene expression

The regulatory activities of TFs were estimated from gene expression data by applying the Weighted Mean method of TF-targets of regulons (confidence levels A-C) provided by decoupleR v2.0.1(62). TF activities per sample were summarized into a statistic corresponding to a normalized weighted mean, and the 50 most variable TFs across samples were selected by their standard deviation for subsequent analyses.

### Regulatory network of TFs modulated by PRMTs

Both experimental and public datasets of TFs with methylation sites regulated by PRMTs and their target genes were handled to outline reported interactions underlying cell-adhesion processes. Proteins with methylation sites regulated by PRMTs were collected from the processed methylomes of PRMT4, PRMT5 and PRMT7(38). Among all the collected PRMTs proteins, only those listed in top 50 TFs with highest variation in estimated activities described above were selected for network building. Additionally, TFs were classified according to the occurrence of arginine methylation sites in their amino acid sequence using public data from dbPTM database (https://awi.cuhk.edu.cn/dbPTM).

Regulatory interactions between the selected TFs and their target genes were established according to the occurrence of TF binding sites within the promoter of genes annotated to cell adhesion from ChIP-Seq data of ReMap2022 catalog(60). Once all the interactions were retrieved (PRMT – TF – adhesion gene), the gene network was built and represented using Cytoscape v3.9.1(63).

### Statistical analysis

The results are expressed as the mean value ± SEM of 1–9 independent experiments. Statistical analyses were performed using ordinary t-Student tests, Mann-Whitney, one-way ANOVA or two-way ANOVA multiple comparisons test depending on the experiments (*p* value <0.05 was considered as significant). GraphPad Prism version 8.4.2 for MacOS X, GraphPad Software, (San Diego, California USA, www.graphpad.com) and R statistical environment(56) were used to represent results. Gene ontology (GO) enrichment of CRISPR/Cas9 library screenings was performed using Metascape(24), gene set enrichment analysis (GSEA), and Human Molecular Signatures Database (MSigDB) C5 as background(64). Venn diagrams were computed using Venny 2.1.0(65).

### Study approval

Slides of FFPE-embedded primary tumor PCa samples were obtained from the Hospital Clínico San Carlos Hospital Biobank (Madrid, Spain) in accordance with protocols approved by the Institutional Review Board. Surgical resection of the primary tumor (PT) from patients with or without de novo metastatic disease were recruited by oncologists Dr. Puente and Dr. Vidal and analyzed by the anatomical pathology unit from Clínico San Carlos Hospital (Madrid, Spain). RNA was extracted following manufacturer’s instructions (High Pure FFPE RNA Micro Kit, Roche). RNA samples with less than 0,01 μg/μL of RNA and less than 0,1 of 260/230 absorbance ratio were excluded from further analysis. Levels of *PRMT7* gene expression between the “High” and “Low” groups were assessed by RT-qPCR using specific primers listed in Supplementary Table 1 at Genetic and Genomic facility of UCM. Gene fold changes were determined by the 2^-ΔΔ^Ct method. GAPDH was used as the reference gene to normalize gene expression.

## Author contributions

Conceptualization: AGU, PB, MRF

Methodology: MRF, AVN, NV, JP, MSP, ALG, MME, NP, CB, AMC, HQ, HH, MM, ARP

Investigation: MRF, AVN, ALG, MM, HQ, ARP, PB, AGU

Visualization: MRF, AVN, MME, HQ, ARP, AGU

Funding acquisition: AGU, PB, AP, ARP, MM

Project administration: AGU

Supervision: AGU, PB

Writing – original draft: MRF, AGU, PB

Writing – review and editing: MRF, AVN, MM, ARP, AP, PB, and AGU

## Acknowledgments

We would like to acknowledge the computing resources and technical support provided by the SCBI (Supercomputing and Bioinformatics) center of the University of Málaga in the Red Española de Supercomputación (RES).

## Data and material availability

All data are available in the main text or supplementary materials.

