## Supplementary material for "CRISPR/Cas9 screenings unearth protein arginine methyltransferase 7 as a novel driver of metastasis in prostate cancer": 309712_0_unknown_upload_8879955_rxy05g_sc.pdf

### Supplemental information

The supplemental information contain supplemental methods, figures and uncropped images of the Western blot results presented in this manuscript.

**1- Supplementary methods.**

**2- Graphical abstract.**

**3- Supplementary figures 1-7.**

### Supplementary methods

#### CRISPR/Cas9 library screening viral production

Lenti-Cas9 viral production was performed in HEK293T cells. A mixture of the 3 transfection plasmids (Packaging and enveloping plasmids psPAX2 (addgene#12260) and pMD2.VSVg (addgene#8454) together with lentiCas9-Blast plasmid (addgene#52962) was prepared and mixed with lipofectamine 2000 (Invitrogen) in a 1:3 ratio and transferred to the HEK293T cells. After 72h, the medium was collected, centrifuged at 2000 rpm for 5 min, and filtered with a 0.45 mm pore syringe filter. PC3 and DU145 cells were transduced with lenti-Cas9 virus to induce the expression of Cas9 in presence of 8 mg/mL of polybrene. 48h later, blasticidine selection (10 mg/ml) was performed for 5 days. Individual clones were expanded and the Cas9 presence was verified by western blot using Cas9 anti-mouse IgG (14697S, CST) and proliferation and migration tests were performed to ensure no clonal effect (data not shown). Subsequently, a mixture of the 3 transfection plasmids (Packaging and enveloping plasmids psPAX2 (addgene#12260) and pMD2.VSVg (addgene#8454) together with the sgRNA from library A (targeting 20,000 genes, including 1,864 sgRNA microRNA and 1,000 non-targeting controls) was prepared and mixed with lipofectamine 2000 in a 1:3 ratio and transferred to the HEK293T cells. After 72h, the medium was collected, centrifuged at 2000 rpm for 5 min and filtered with a 0.45 mm pore syringe filter.

#### Exogenous PRMT7 transfection to cells

For exogenous PRMT7 expression pCDH1-PRMT7-GFP plasmid was obtained from addgene[37]. 4 ug of the plasmid were transfected to PRMT7 depleted and control PC3 using lipofectamine 2000 in a 1:3 ratio. Subsequently, proteins were extracted and levels of ITGb4 and ITGa1 were analyzed as described in Materials and methods of this study.

### Graphical abstract

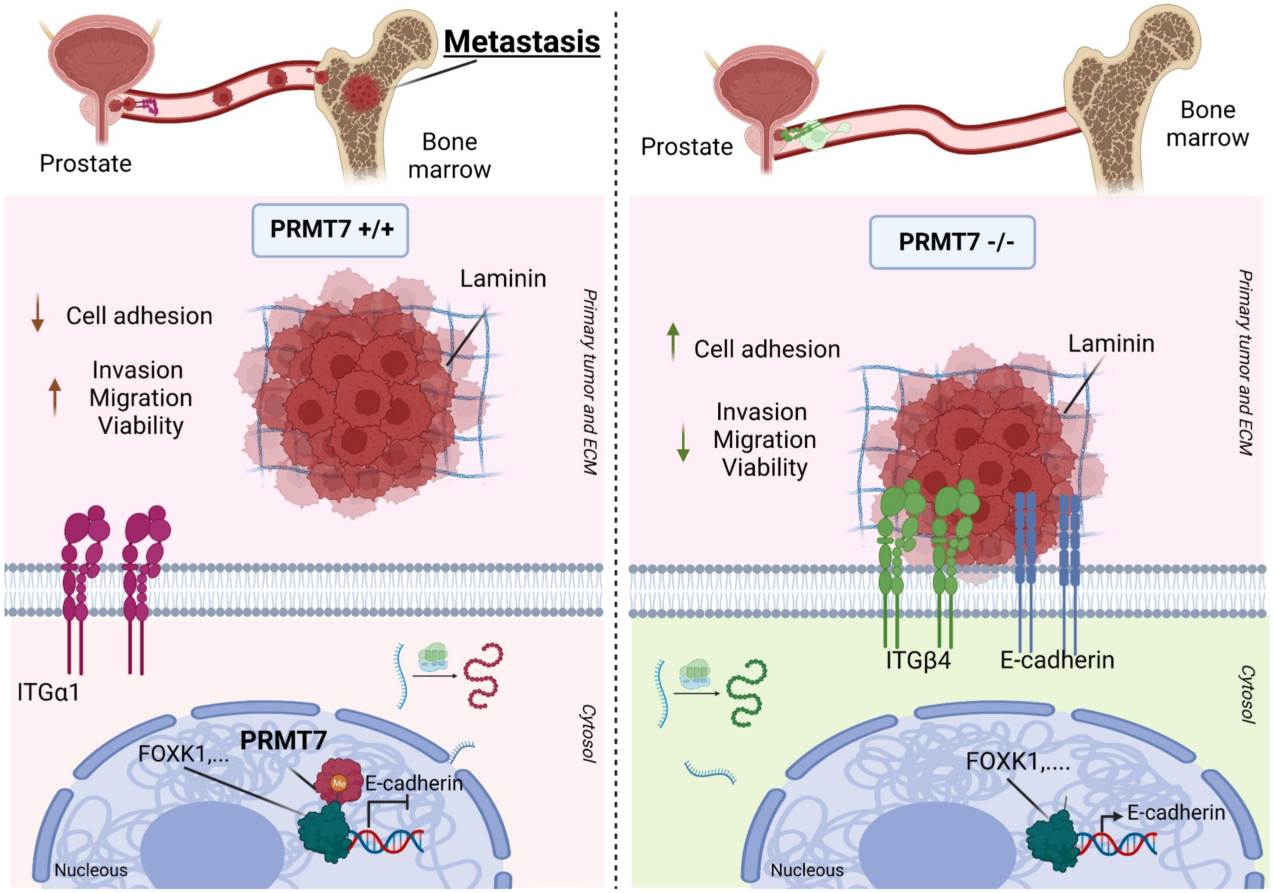


**Graphical abstract. Schematic representation of the relevance of PRMT7 in prostate cancer metastasis (mPCa) onset.** *PRMT7*, identified as a novel metastasis regulator by CRISPR/Cas9 high-throughput screening in two different mPCa cell lines, PC3 and DU145, was validated *in vitro*, *in ovo* and *in vivo*. *PRMT7* depletion significantly reduces migratory, invasive and proliferative capacities of cells *in vitro* and *in vivo*. However, its depletion increases cell adhesion by producing a switch of the cell-surface adhesion molecules expressed by cancer cells, that was uncovered by our differential transcriptomic analyses. Altogether, P*RMT7* genetical inhibition or depletion or pharmacological inhibition reduces the primary tumor dissemination, pointing it out as a novel important mediator of mPCa onset and as a promising mPCa prevention target.

### Supplementary Figure 1

**A**

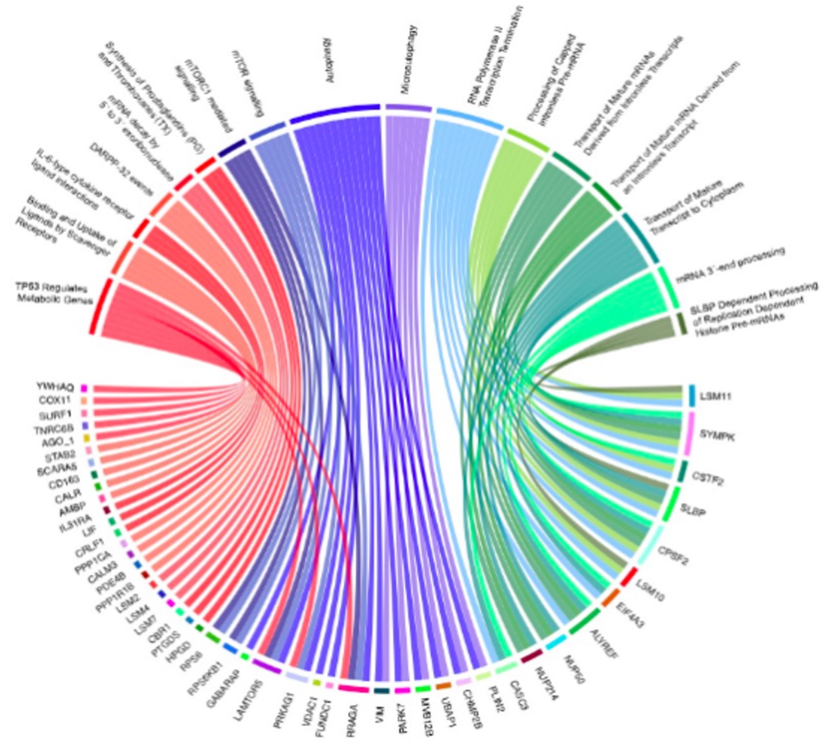


**B**

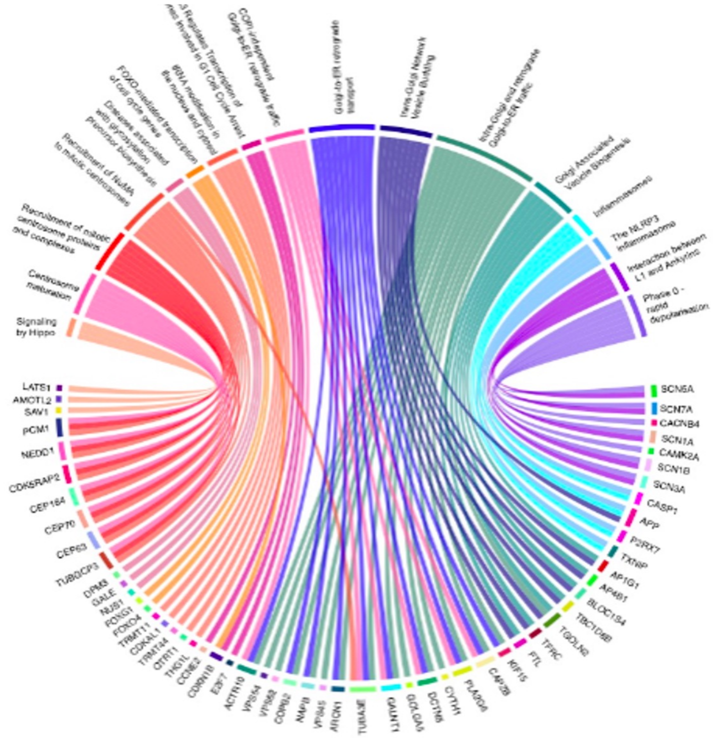


**Supplementary Figure 1. Gene set enrichment analysis (GSEA) using Reactome database. A-B** Results of GSEA Reactome analysis of **A,** PC3, **B,** DU145 screening results.

### Supplementary Figure 2

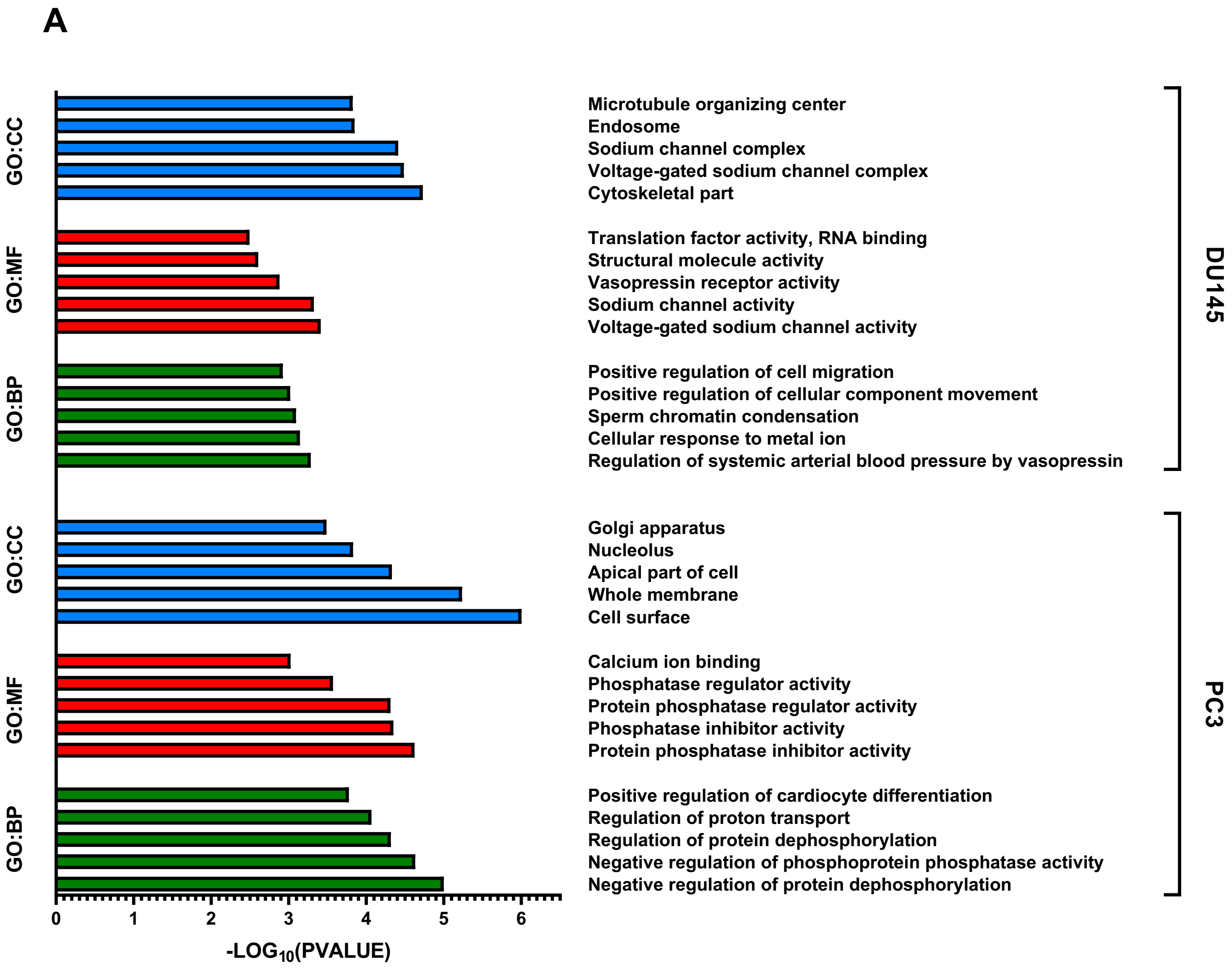


**Supplementary Figure 2. Gene ontology enrichment analyses. A,** Graphical representation of gene ontology enrichment analysis of gene ontology cellular component (GO:CC), gene ontology molecular function (GO:MF) and gene ontology biological pathway (GO:BP) analyses, for DU145 and PC3, respectively.

### Supplementary Figure 3

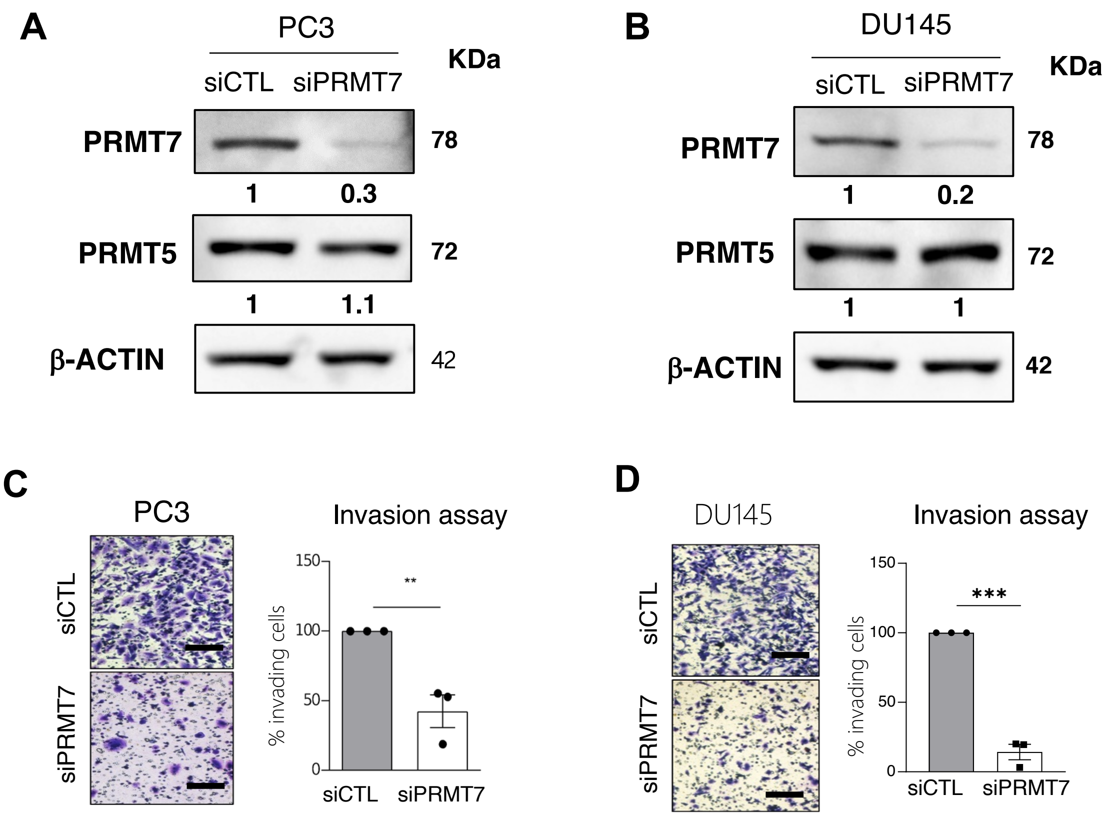


**Supplementary Figure 3. Analysis of the effect of *PRMT7* inhibition by siRNA in PC3 and DU145 cells. A-B,** Western blot analysis showing a reduction of PRMT7 expression levels while PRMT5 expression levels remain unaltered, for **A,** PC3, **B,** DU145 cells. **C-D,** Invasion assay using FBS as chemoattractant of siPRMT7 inhibited cells versus siCTL cells of **C,** PC3 and **D,** cell lines.

### Supplementary Figure 4

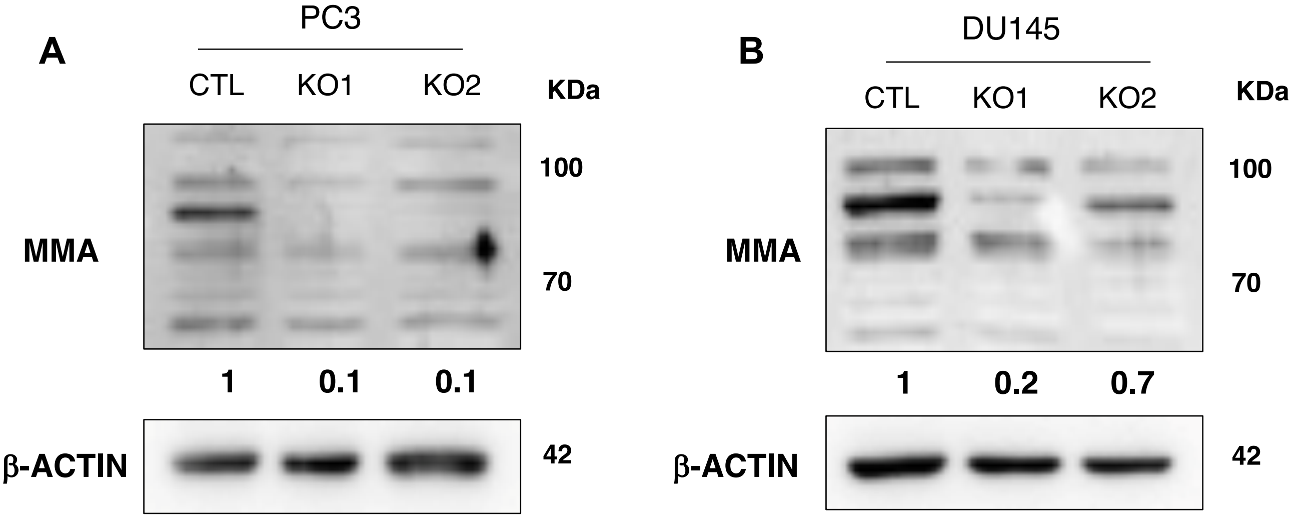


**Supplementary Figure 4. PRMT7 depleted cells show reduced levels of MMA and lower viability. A-B,** Western blot analyses of MMA levels upon PRMT7 genetic depletion in **A,** PC3-Cas9 and **B,** DU145-Cas9 cells.

### Supplementary Figure 5

A

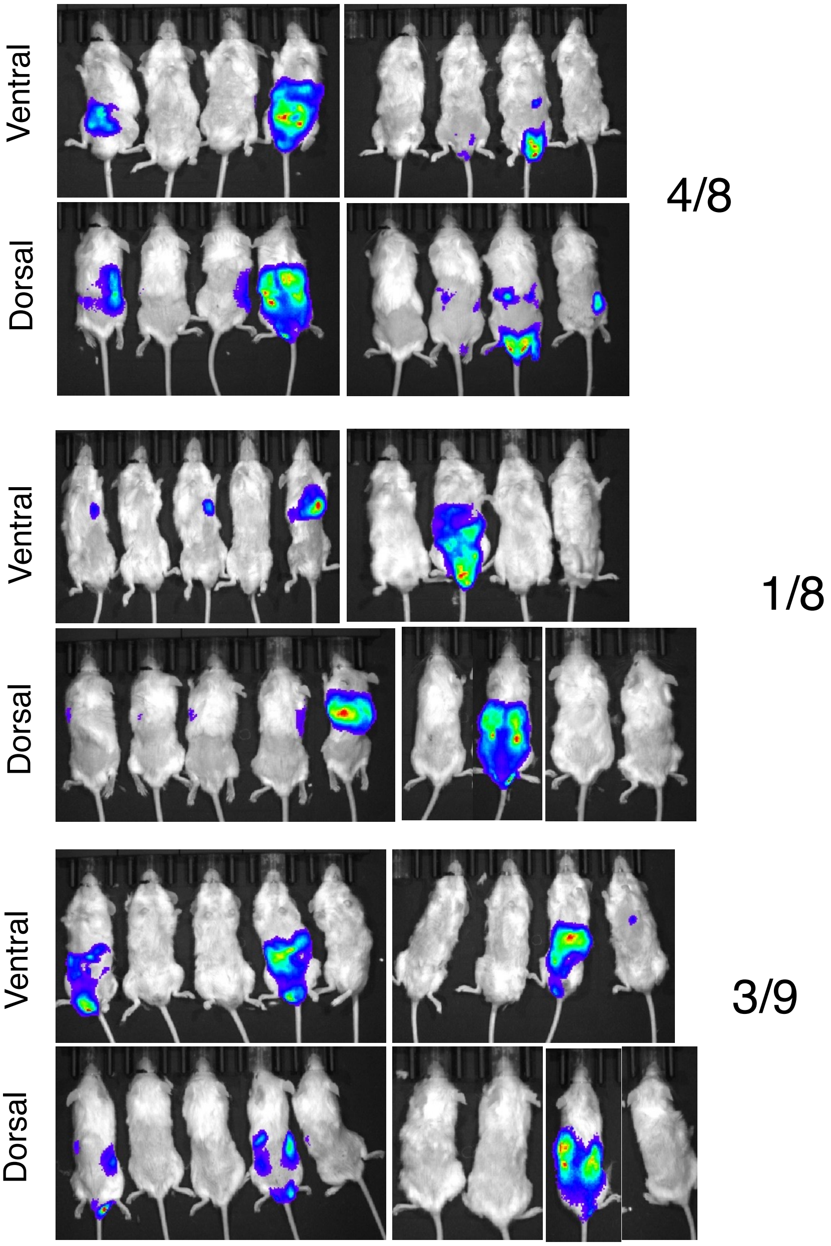


B

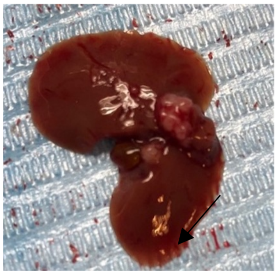

**Supplementary Figure 5. Macrometastasis observed upon luciferine inoculation in mice A,** Picture of mice in the IVIS imager **B,** Picture of liver macrometastasis in a mouse inoculated with CTL PC3 cells.

### Supplementary Figure 6

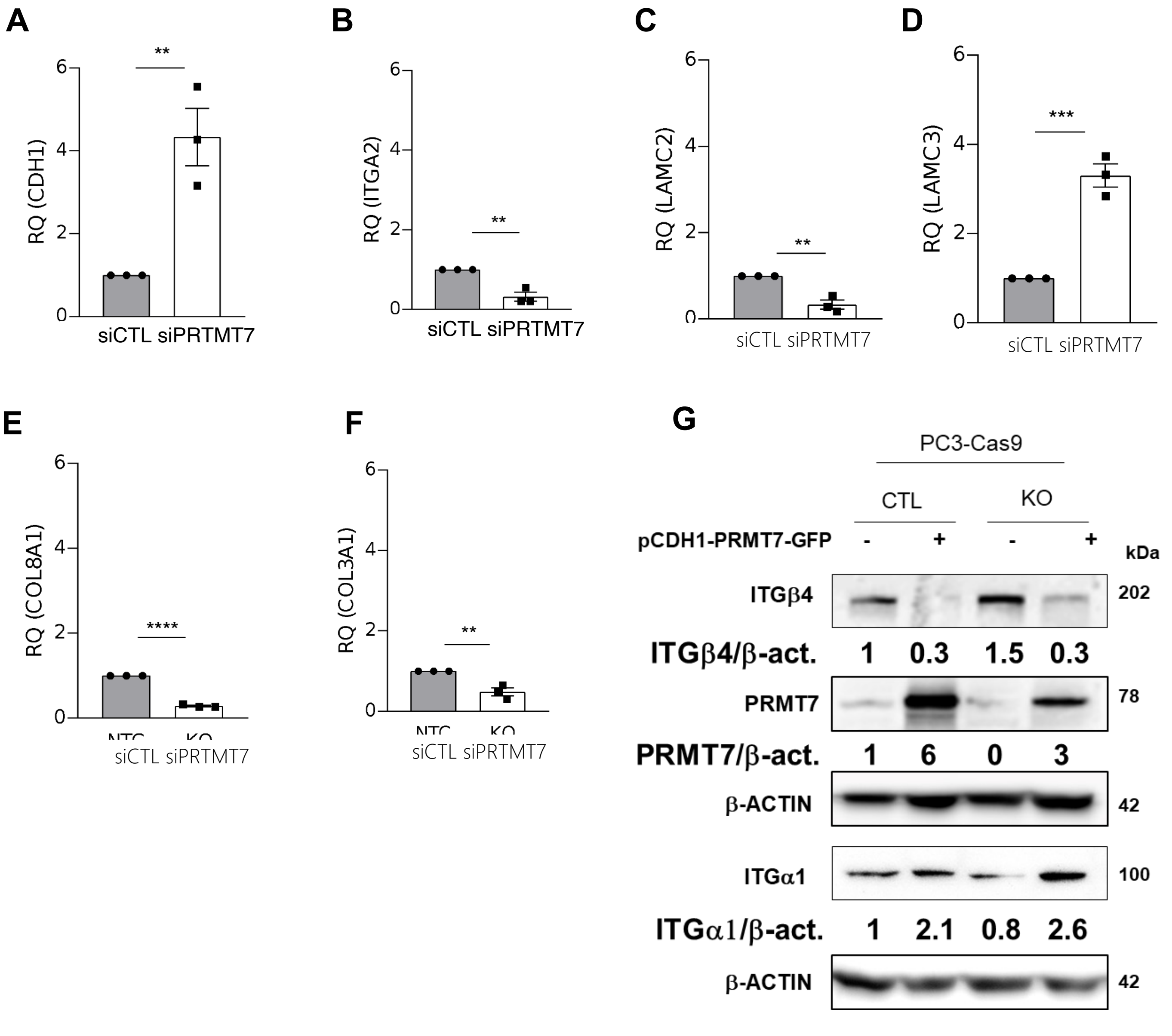


**Supplementary Figure 6. PRMT7 produces a cell adhesion molecules gene expression switch in PC3 cells. A-F** qPCR validation of **A** *CDH1*, **B** *ITGA2*, **C** *LAMC2*, **D** *LAMC3*, **E** *COL8A1*, **F** *COL3A1,* **G** Western blot showing ITGα1 and ITGβ4 protein levels upon exogenous PRMT7 overexpression in PC3 cells.

### Supplementary Figure 7

**A**

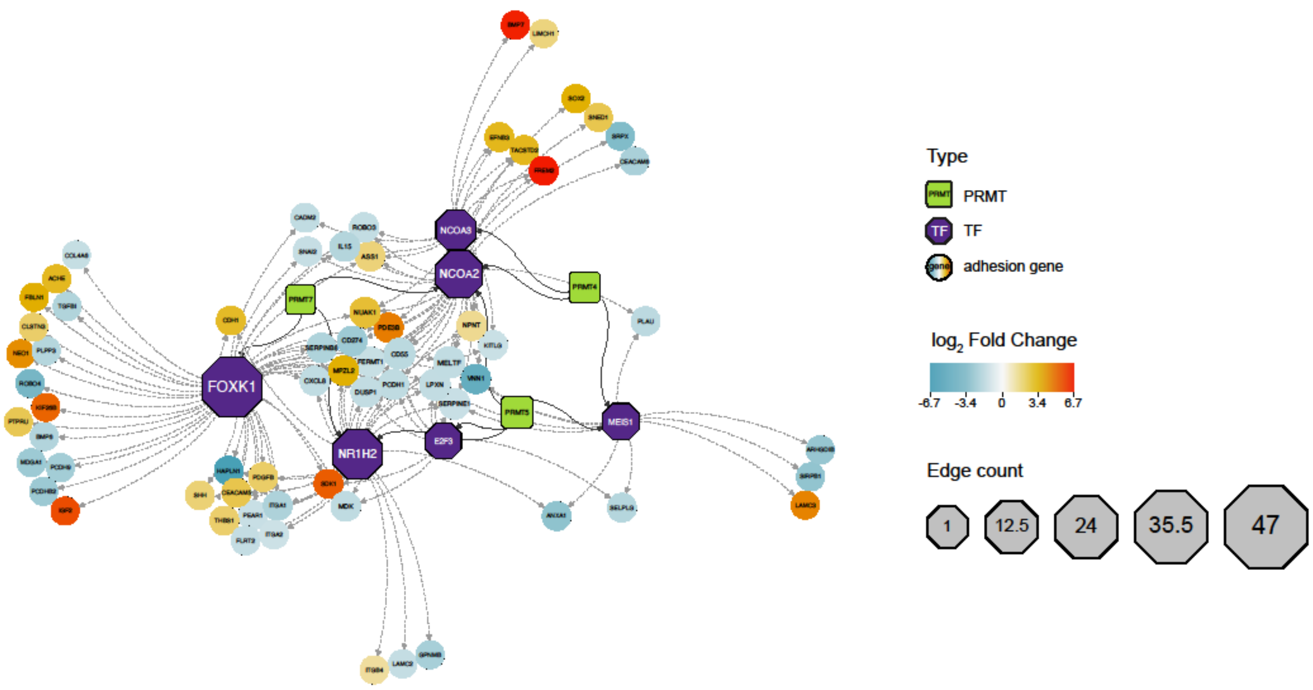


**Supplementary Figure 7. PRMT4, PRMT5 and PRMT7 transcription factor regulomes.**

**A,** Regulatory network of the three PRMTs (green squares) that methylate transcription factors (purple hexagons), and that bind to promoters of cell adhesion genes (circles) that are differentially expressed. Fold changes calculated in the differential gene expression analysis (PRMT7 depleted cells compared to control cells) are represented by color. Edge count (size) indicates the number of connections per item of the network.

### Uncroppeds of Figure 3B

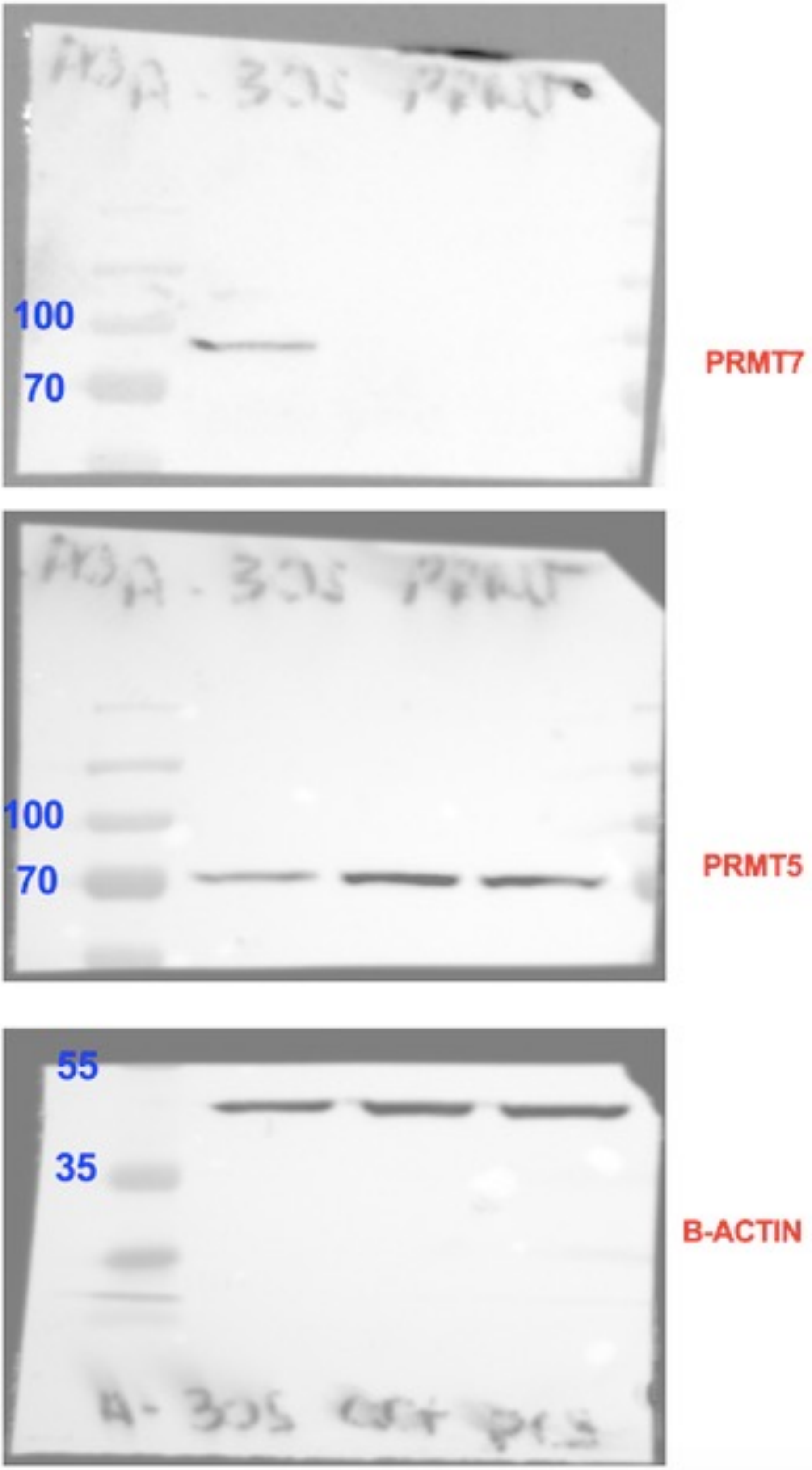

### Uncroppeds of Figure 3C

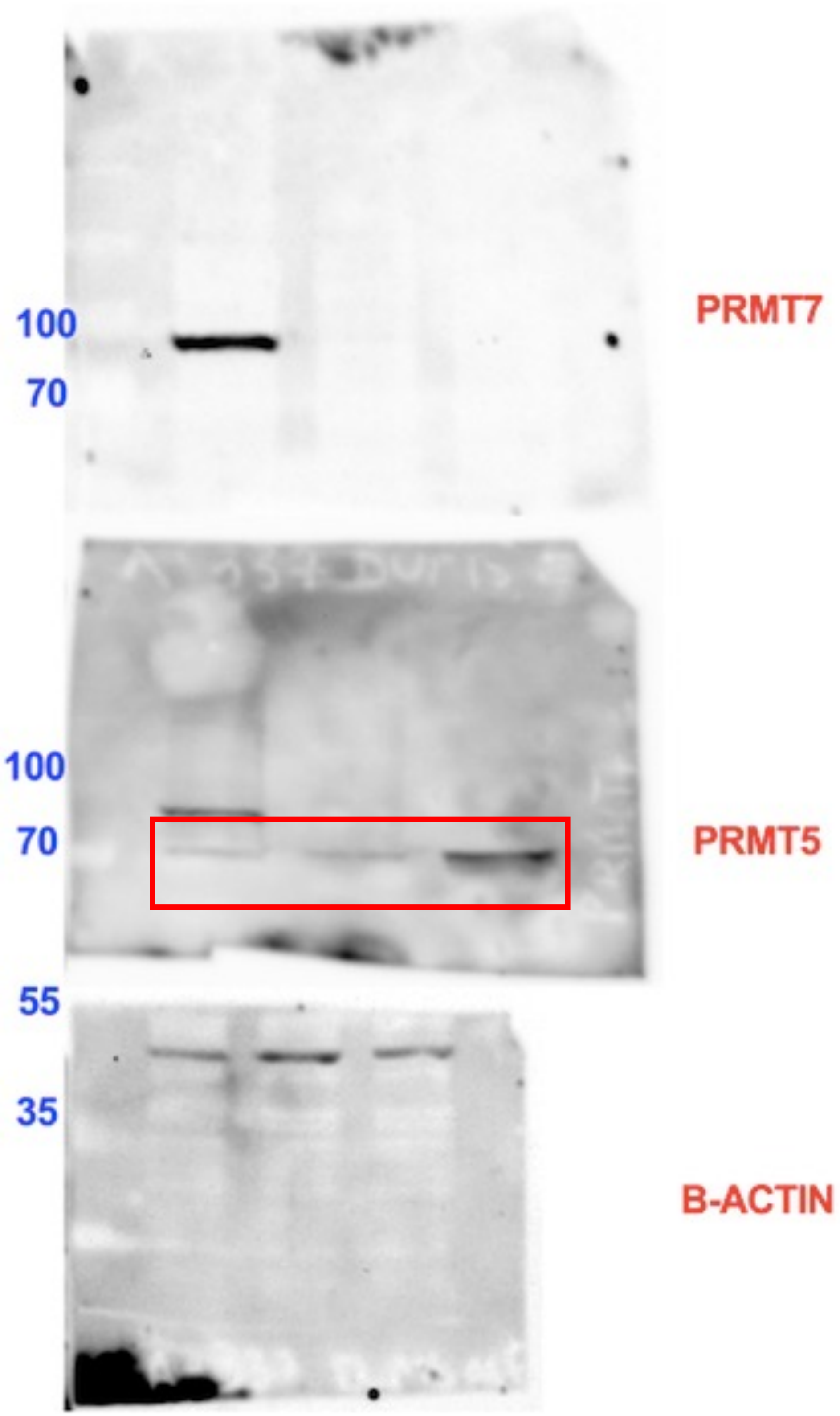

### Uncroppeds of Figure 4A

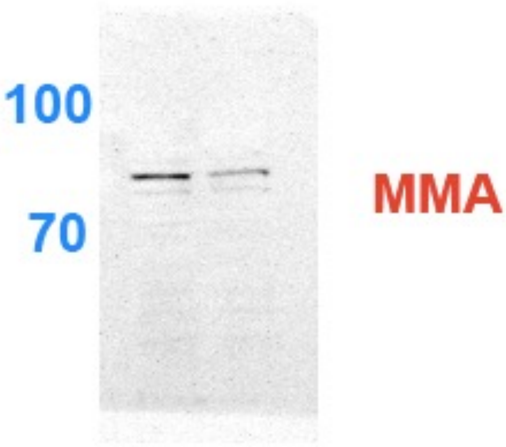


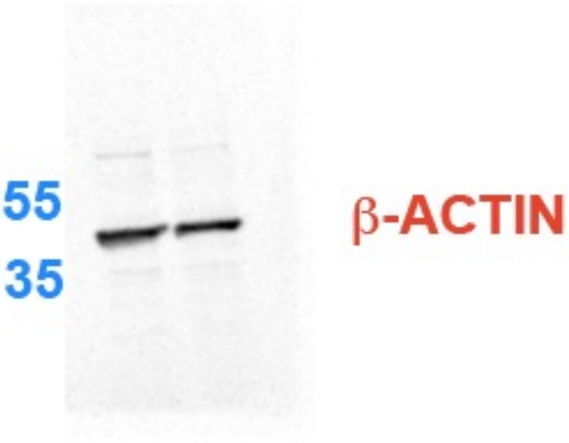

### Uncroppeds of Figure 4B

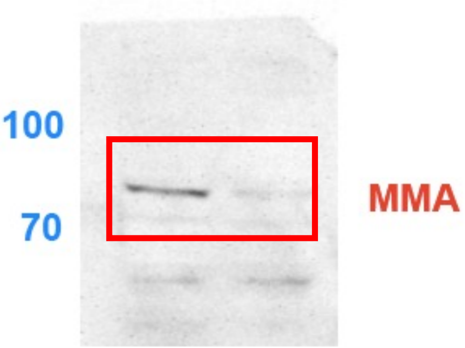


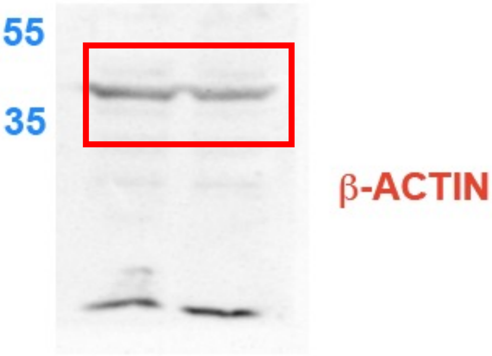

### Uncroppeds of Figure 6D

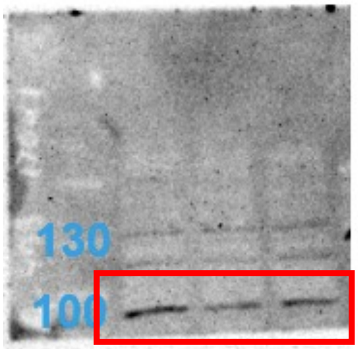


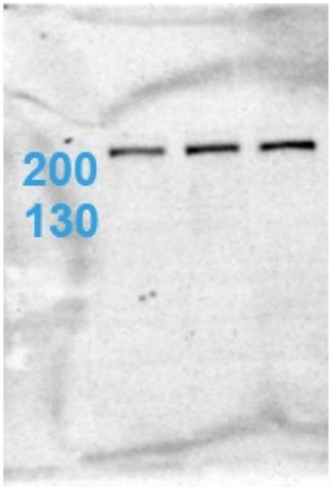


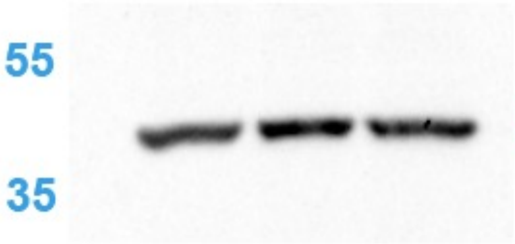

### Uncroppeds of Figure 6E

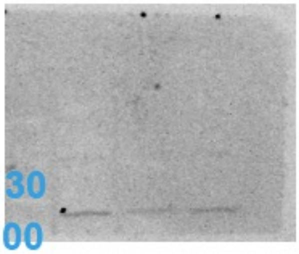


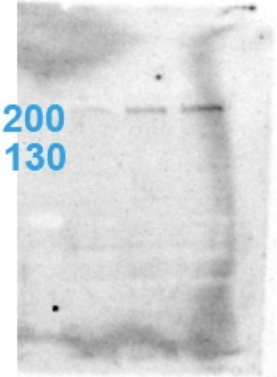


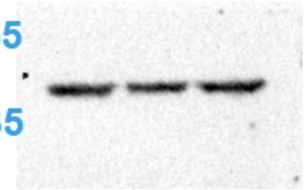

### Uncroppeds of Supplementary Figure 2A

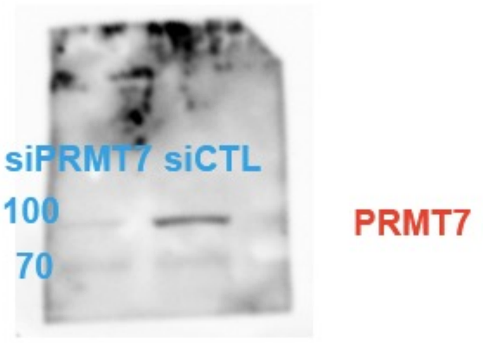


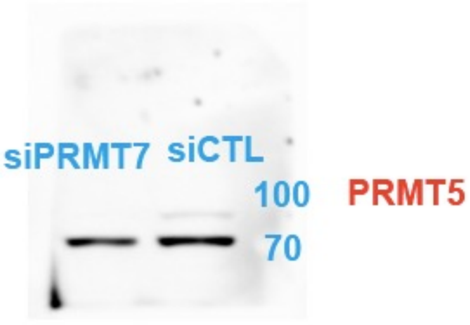


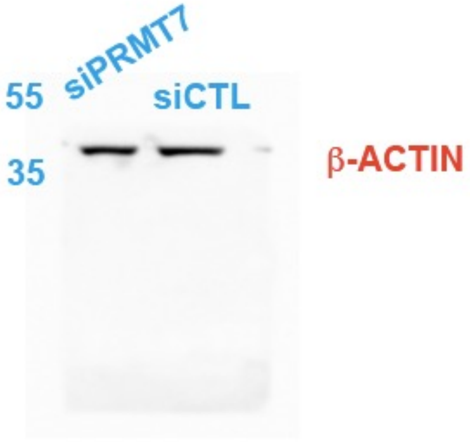

### Uncroppeds of Supplementary Figure 2B

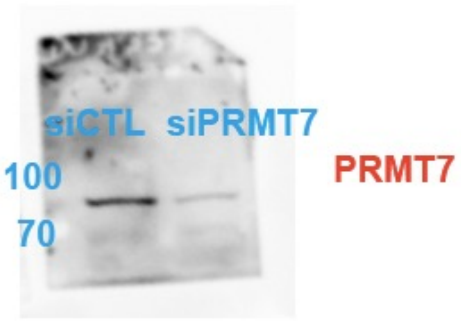


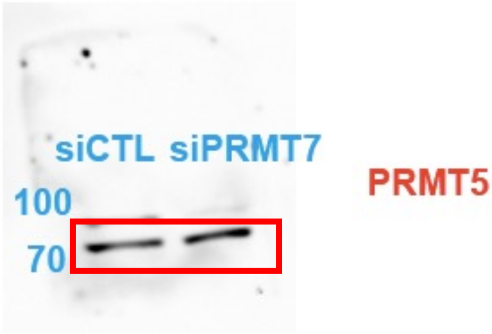


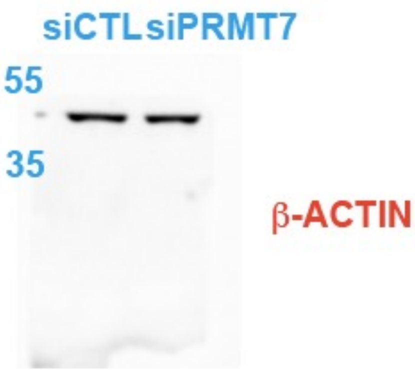

### Uncroppeds of Supplementary Figure 3A

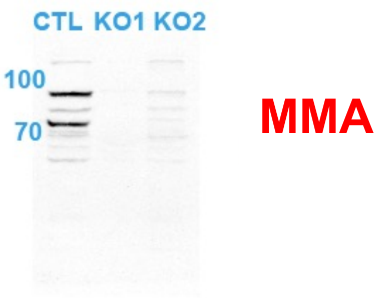


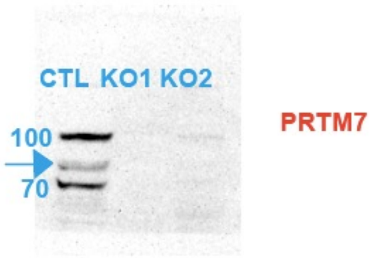


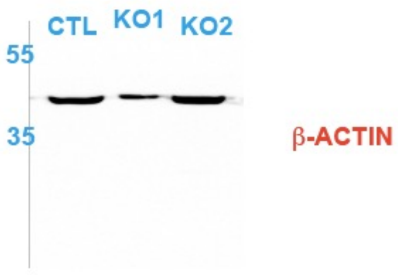

### Uncroppeds of Supplementary Figure 3B

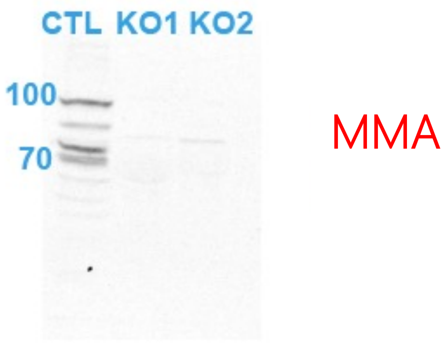


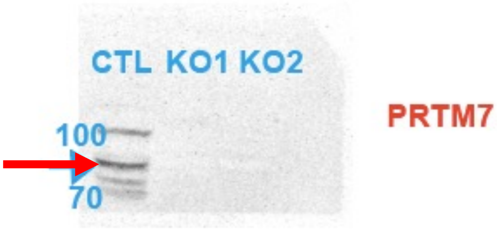


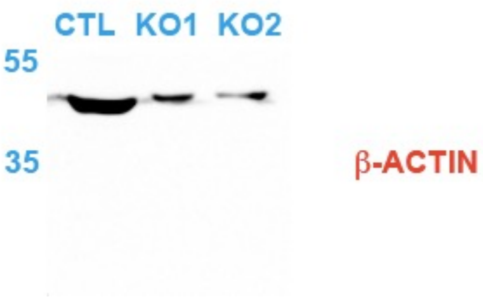

### Uncroppeds of Supplementary Figure 6G

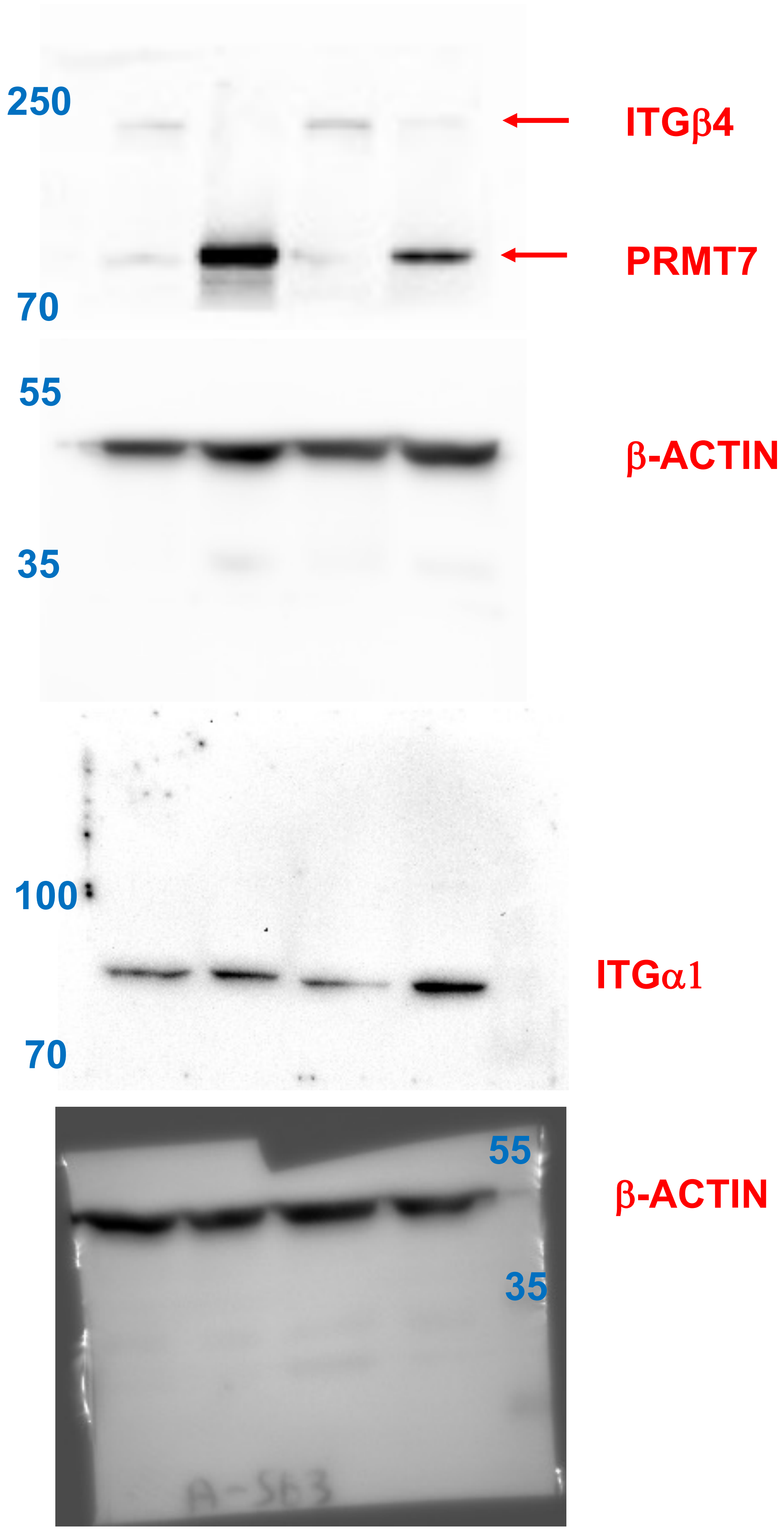
